# Aortic annuloplasty FSI digital twin of 3D-printed phantoms with 4D-flow MRI comparison

**DOI:** 10.1101/2025.03.05.641637

**Authors:** Luca Bontempi, Marta Zattoni, Anna Ramella, Francesco Migliavacca, Steffen Ringgaard, Won Yong Kim, Peter Johansen, Monika Colombo

## Abstract

**Background:** Aortic annuloplasty, involving the implantation of an external ring around the aortic root to reduce annular dimensions, is a promising treatment for aortic valve insufficiency. However, its hemodynamic effects remain underexplored due to the absence of computational models validated by experimental and clinical data.

**Methods:** This study introduces a computational fluid-structure interaction (FSI) model of supra valvular aortic annuloplasty using 4D-flow magnetic resonance imaging (MRI). Native and post-annuloplasty conditions of idealized aortic root phantoms, including the aortic valve, were CAD-modelled and 3D-printed with elastic resin. These phantoms were tested in a mock circulatory flow-loop providing normal pulsatile physiologic conditions using a glycerol-water mixture to simulate blood viscosity. Flow and pressure data collected from sensors were used as boundary conditions for FSI simulations. Experimental velocity fields from 4D-flow MRI were compared to computational results to assess model accuracy.

**Results:** MRI scans of the annuloplasty model showed an increased peak systolic velocity (up to 145.4 cm/s) and localized flow alterations, corresponding to a higher pressure gradient across the valve. During regurgitation, the annuloplasty model showed broader velocity distributions compared to the native condition. The FSI simulations closely matched 4D-flow MRI data, with strong correlation coefficients (r > 0.93) and minimal Bland-Altman differences, particularly during systolic phases.

**Conclusions:** This study establishes an integrative methodology combining in-vitro, in-silico, and clinical imaging techniques to evaluate aortic annuloplasty hemodynamics. The validated digital twin framework offers a pathway for patient-specific modelling, enabling prediction of surgical outcomes and optimization of aortic valve repair strategies.

## 1 Introduction

Aortic insufficiency is a condition affecting the aortic valve (AV), often associated with aortic root (AR) dilation [1]. If AR dilation is the primary issue and the AV leaflets remain intact, valve replacement may not be necessary to resolve the condition. Preserving the native valve is essential to mitigate potential complications linked to prosthetic valve replacement [2]. Aortic annuloplasty (AA) emerges as a repairing approach and consists of the implantation of a ring placed externally around the AR, reducing annular dimensions and restoring AV function [3–6]. Recently, a novel open annuloplasty ring [7], has been developed allowing for its positioning without the prior removal of the coronaries, thereby reducing the risk of the surgical procedure [8].

To assess the AA interventional approach, experimental testing systems such as flow-loop test bench [7,9,10] and computational analysis can be effectively combined to understand the effects on the AR hemodynamics [11–13]. Experimental approaches are typically time and cost consuming, with strict assumptions of geometries and testing conditions. Recent FDA guidelines recognize computational model simulations as an alternative method for assessing applicability in the clinical practice and validation of medical devices [14]. However, challenges persist with computational fluid-structure interaction (FSI) algorithms, including the need for realistic representations of fluid and structural domains, computational costs, and validation against experimental or clinical data [15,16]. Notably, no computational model of annuloplasty for the AV has been documented in the literature to date.

This study aims to evaluate the impact and consequences of AA on AV hemodynamics using an idealized AR model. To achieve this, an integrative methodology was developed, incorporating 3D-printed AR phantoms to analyse both native and post-annuloplasty hemodynamic conditions across multiple levels of investigation. Initially, in-vitro mock circulatory flow-loop (MCL) testing campaign [17] was conducted to establish appropriate flow and pressure conditions. Subsequently, an FSI-based digital twin model of AA was compared against 2D and 4D-flow magnetic resonance imaging (MRI) data obtained from the in-vitro MCL experiments.

## 2 Material and methods

### 2.1 Integrated workflow

The methodology employed to integrate measured flow and pressures sensor-based data, in-silico simulations of native and post-operative AA conditions, and clinical imaging data of the experimental setup via a pilot study on AA is outlined in Fig. 1. The computer aided design (CAD) model of idealized ARs was used to create both the 3D-printed phantoms for integration into the MCL and the geometries for computational simulations. The hemodynamic data collected from MCL flow and pressure sensors were incorporated into the in-silico model as boundary conditions to perform FSI simulations. Finally, the fluid velocity field of the in-vitro phantom was assessed through 4D-flow MRI scans, and the comparison with the post-processed FSI results facilitated evaluation of the AA digital twin.

**Fig. 1.**
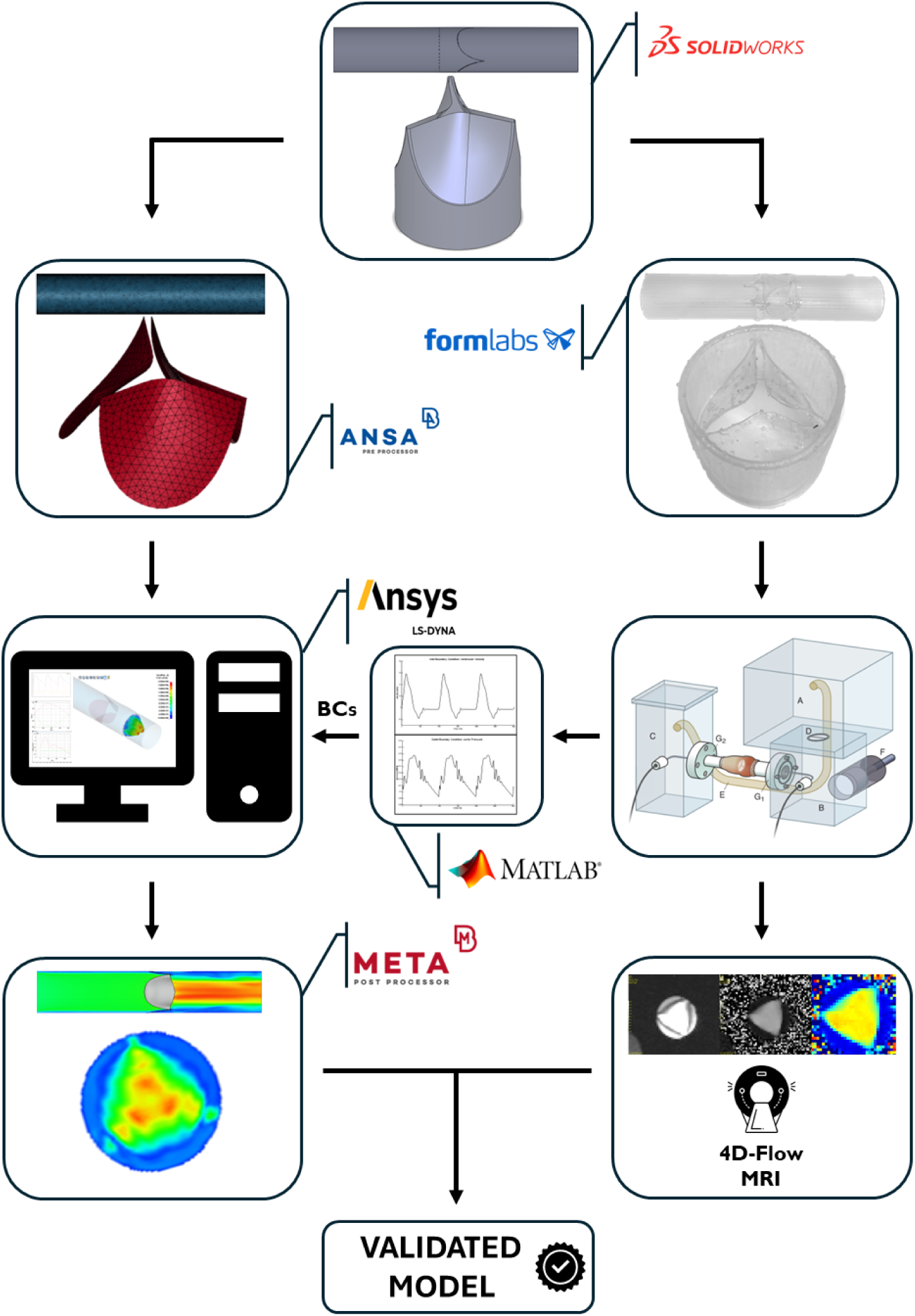
Workflow of the study: CAD model of idealized aortic root used for creating 3D-printed phantoms for mock circulatory flow-loop experiments and defining boundary conditions for FSI simulations. Velocity fields from 2D- and 4D-flow MRI scans were compared with the fluid-structure interaction results to evaluate the aortic annuloplasty digital twin performance.

### 2.2 Aortic annuloplasty model

An idealized AR model was designed omitting the Valsalva sinuses for the sake of simplicity in obtaining a 3D-printed model, with the primary focus on accurately capturing the morphology of the leaflets. A 150-mm long hollow tube with a 27-mm internal diameter was chosen to emulate the aortic wall, as previously done [18,19]. A 1.5-mm thickness was set for the cylindrical structure to achieve a compliance value close to 1 ml/mmHg, similar to the native AR as identified through both an echocardiographic study [20] and an in-vivo investigation [21].

Obtaining a physiological leaflet thickness of 0.1-0.2 mm [22] through 3D-printing was proven unfeasible with the available equipment (see *Supplementary Information* (SI) section S.1 for further details on the 3D-printing protocol). Consequently, a thickness of 0.4 mm was selected and led to a successful printing of the valve. The valve’s geometry was designed to optimize leaflet coaptation, ensuring the most competent valve function possible within the constraints of the material.

To investigate the hemodynamic consequences of the implantation of the AA ring on the native model, a reduction in the cylinder diameter downstream the valve was implemented (Fig. 2). To replicate experimental conditions, the AA ring was implanted in a non-pathological AR model, as preclinical studies lack a model for aortic insufficiency [7]. In accordance with the clinical data [7], a 20% reduction in area was selected to emulate the presence of the external ring. The area constraint was introduced at one diameter downstream the valve to ensure a smooth printing process for the leaflets. To achieve a 20% reduction in area, i.e. from 573 mm^2^ to 458 mm^2^, a final diameter of 24.15 mm was generated to create the restriction downstream the valve; the reduction was applied over a 4-mm length of the tube to align with the ring height [7].

**Fig. 2.**
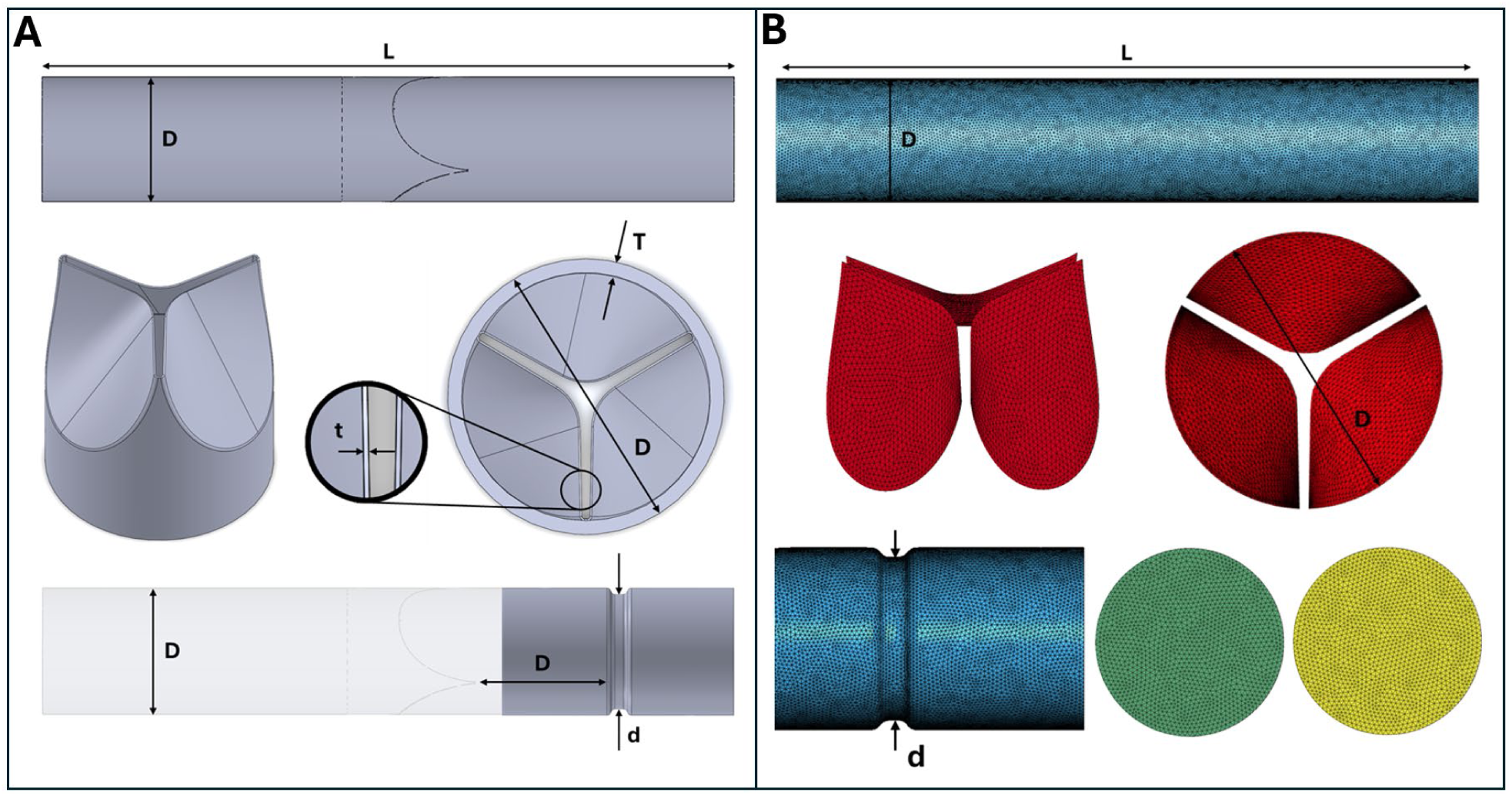
A) Native (top) and annuloplasty (bottom) aortic root CAD model employed for the generation through 3D-printing of the phantoms and the in-silico model geometries. L: length (15 cm); D: internal diameter (27 mm); T: wall thickness (1.5 mm); d: annuloplasty internal diameter (24.15 mm); t: leaflet’s thickness (0.4 mm). **B)** Detail of the final computational mesh of the native (top) and annuloplasty (bottom) models.

All the employed samples underwent the same 3D-printing protocol (see SI section S.1). Initially, CAD models were crafted with SolidWorks 2022 (Dassault Systèmes SolidWorks Corp., Waltham, MA, USA), followed by the exportation of an .STL file to Preform (Formlabs Inc., Somerville, MA, USA), where orientation and support placement were specified. A flexible resin, Elastic 50A (Formlabs Inc., Somerville, MA, USA) was employed in the Form 3B+ printer in the MAKERLab Pro facility (Aarhus University, Department of Electrical and Computer Engineering), given its similarity to natural aorta mechanical properties, as reported by the company [23] and verified through mechanical characterization (see SI section S.2 for further details). After printing, the samples were rinsed in isopropyl alcohol for 20 minutes and cured under UV light at 60°C for an additional 20 minutes, following the manufacturer’s postprocessing guidelines.

### 2.3 In-vitro flow measurements

A pulsatile in-vitro MCL was employed to replicate a left heart flow cycle, as previously described [9,10]. A glycerol/demineralized water mixture in the volume ratio 1:3 was used as test fluid to obtain blood-like fluid properties [24]. Fig. 3 shows the components of the flow-loop, namely a left atrium reservoir, a left ventricle chamber, the 3D-printed AR phantom, an air-controlled compliance chamber, and a peripheral resistance clamp. Both compliance and peripheral resistance were adjustable within the system. An electromechanical piston pump (VSI Superpump, ViVitro Labs, Victoria, Canada) was connected to the ventricular chamber to generate pulsatile flow and pressure waveforms. The atrial reservoir and ventricular chamber were linked via a mechanical heart valve mimicking the mitral valve, while the ventricular chamber and the compliance chamber were connected through the AR model with the integrated AV.

**Fig. 3.**
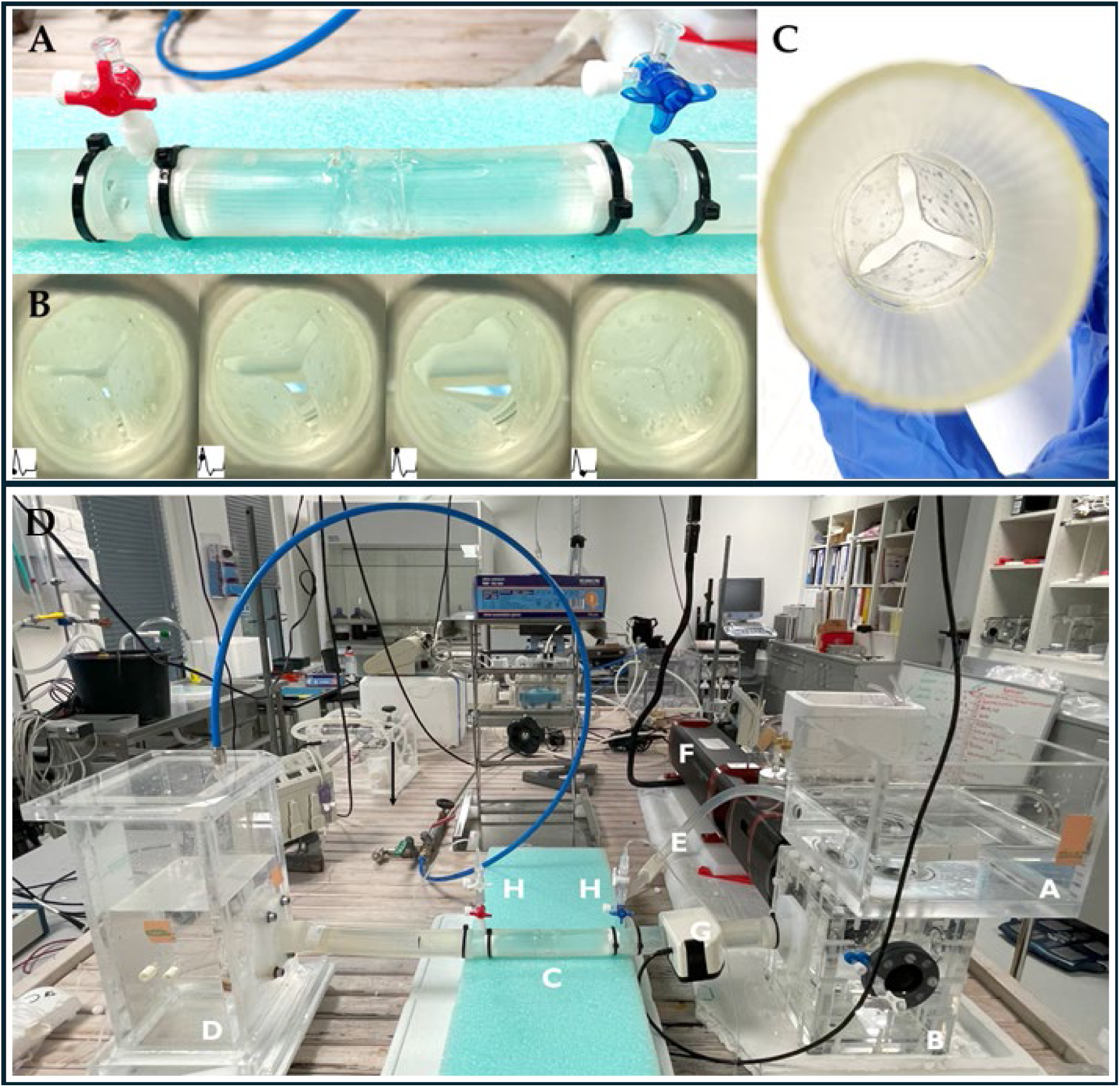
**A**) 3D-printed AR phantom connected to two rigid connectors for pressure readings; **B**) short-axis view of the 3D-printed valve during four different phases of the cardiac cycle; **C**) 3D-printed AR phantom after UV curing and supports removal; **D**) CAVE Lab mock circulatory flow-loop with the AR phantom connected (in white, A – atrial chamber, B – ventricular chamber, C – AR phantom, D – compliance chamber, - E – systemic resistance, F – pulsatile pump, G – flowmeter, H – pressure sensors).

The flow through the valve was monitored using a transit time flowmeter (ME-20PXL, Transonic Systems Inc., Ithaca, NY, USA) placed right upstream the AV. Continuous pressure recordings both upstream (ventricular pressure) and downstream (aortic pressure) the AV were achieved using micro-tip pressure catheters (SPC-350, Millar Inc., Houston, TX, USA) and a signal conditioning unit (Millar Inc., Houston, TX, USA). To precisely capture the pressure curves near the AV, custom 1-inch connectors were designed and fabricated through 3D-printing, utilizing rigid resin. These connectors serve as crucial components, facilitating the connection of the model to tubes linked to the ventricular and compliance chambers. Simultaneous acquisition of flow and pressure data at a sampling rate of 1000 Hz was accomplished using a multipurpose 16-bit data acquisition module (NI USB-6259, National Instruments, Austin, TX, USA).

Measurements from the MCL were obtained by converting recorded voltage signals into pressure and flow values (see SI Fig. S3) using a custom LabVIEW virtual instrument (National Instruments Inc., Austin, TX), calibrated to transducer sensitivities. Flow and pressure data were initially recorded using LabVIEW, and further analysed in MATLAB (The MathWorks Inc., Natick, Massachusetts, United States). This analysis involved computing flowrate, ventricular and aortic pressures, as furtherly detailed in SI section S.3.

### 2.4 Fluid-structure interaction model

#### 2.4.1 Meshing and pre-processing

The native and annuloplasty AR models were realised with ANSA pre-processor (v23.1.3, BETA CAE Systems, Switzerland). The geometry was discretized beginning with the AV. Given that the thickness-to-width ratio was below 0.1, the leaflets were considered thin structures and discretised, after reaching a grid convergence (reported in SI section S.4 and Fig. S4.1-2), with 5,740 3-node shell elements (Fig. 2-B). After mirroring the commissure lines’ nodes on the tube to maintain consistent node and element IDs between the two parts, the cylindrical structure was also discretised with 65,607 3-node shell elements. Two surfaces were generated adjacent to the proximal (inlet) and distal (outlet) ends of the tube, creating a closed volume, and discretised with 2,941 and 2,884 3-node shell elements, respectively. The structural part comprised a total of 77,172 triangular shell elements. The fluid volume mesh, consisting of 1.9 million tetrahedral Eulerian elements with a single iteration point and characteristic dimension of 0.65 mm, was automatically generated by the finite element solver from the closed volume defined by the structural parts. To manage computational costs, linear elements with reduced integration were favoured.

The same discretization process was adopted for the structural mesh of the annuloplasty model, by keeping the same element characteristic dimension of 0.65 mm. The obtained mesh for the annuloplasty model comprised 79,361 3-node shell elements, where a total of 67,592 elements was used to discretize the tube (+3% with respect to the native model). Finally, 2,05 million tetrahedral Eulerian elements with a single iteration point were generated, as shown in the detail in Fig. 2-B.

A summary of the number of structural and fluid elements used for both models is provided in SI section S.4.

#### 2.4.2 Numerical simulations

Simulations were conducted using the commercial finite element solver LS-Dyna 971 R14.0 (Ansys, Canonsburg, PA, USA). A strong two-way FSI coupling method was selected, and the full non-linear problem was solved using an implicit solver for the Lagrangian mesh. Both the native and annuloplasty AR models were modelled using a linear elastic material, and the material properties for the 50A resin were derived from the material characterization conducted at the MaterialsLab facility (Aarhus University, Department of Mechanical and Production Engineering), reported in SI section S.2. Both the leaflets and the cylindrical structure were assigned a density of 1100 kg/m^3^, a Young’s modulus of 1.99 MPa, and a Poisson’s ratio of 0.45. The fluid, i.e., the glycerol/water mixture, was modelled as a Newtonian fluid with a density of 1050 kg/m^3^ and a dynamic viscosity of 0.0035 Pa·s [24], accordingly to experimental conditions. A no-slip boundary condition was applied on the fluid nodes located at the internal surface of the AR.

The final FSI simulations consisted of three complete cardiac cycles by applying at the inlet experimental physiological time-dependent ventricular velocity and at the outlet the experimental aortic pressure. Only the results from the third cardiac cycle were analysed to ensure numerical robustness. Throughout the entire simulation, the nodes on the proximal and distal ends of the cylindrical structure were constrained in terms of translations and rotations.

Aortic leaflets coaptation was simulated using the automatic scale-penalty contact algorithm provided in LS-Dyna, activating the soft constraint option: during diastole, automatic single surface contact was defined between the leaflets, with friction coefficient of 0.05 [25], viscous damping coefficient of 50, scale factor for penalty stiffness of 0.5, for contact thickness of 1.0, for sliding interface penalties of 0.1. The choice of these parameters was based on their impact on simulation accuracy, with an emphasis on maintaining consistent contact forces and reducing numerical issues, such as penetration between components [25].

The selection of these parameters was based on their impact on simulation accuracy, considering the consistency of contact forces and the reduction of numerical artifacts, such as penetration between different parts.

Similarly to previous works, a mass proportional damping factor of 10 was applied to the leaflets, to control excessive vibrations, ensuring that the ratio of dissipated to internal energy and of kinetic to internal energy remained below 5% during the simulation [25,26].

Time discretization was automatically set to default values by dynamically satisfying the Courant-Friedrichs-Lewy (CFL) condition, optimizing numerical stability throughout the simulation.

The Lagrangian time integration step was initially set to 0.5 ms, with limits ranging from 0.1 to 1 ms, and was calculated using the central difference method to ensure second-order accuracy. For the nonlinear implicit analysis, the Newmark method was employed, with time integration constants *gamma* and *beta* set to 0.6 and 0.4, respectively.

The total computation time for the simulation, employing the optimal mesh and performed on 28 CPUs of an Intel Xeon64 system with 250 GB of RAM, was approximately 16 hours. Post-processing analysis was carried out with META post-processor (v23.1.3, BETA CAE Systems, Switzerland) and ParaView 5.11.1 (Sandia National Labs, Kitware Inc., Los Alamos National Labs) for results’ visualization and quantitative analysis.

### 2.5 4D-flow MRI: acquisition settings and post-processing

Phase-contrast 2D-flow and 4D-flow MRI scans ensured a direct visualization and comparison of the fluid dynamics patterns in terms of velocity components in both the native and annuloplasty models. Images were acquired using a clinical 1.5 T scanner (MAGNETOM Sola, Siemens Healthineers, Erlangen, Germany), located at the Department of Cardiology, Aarhus University Hospital, Denmark. The velocity encoding (Venc) was consistently maintained at 170 cm/s to ensure comprehensive capture of the fluid velocity magnitude [28]. Notably, 2D-flow scans, performed in both short and long axes with a resolution of 1.1 x 1.1 x 5 mm, were completed within approximately 4 minutes each. Conversely, 4D-flow scans were four times longer, totalling approximately 16 minutes, albeit at a slightly reduced spatial resolution (1.9 x 1.9 x 1.9 mm). To enable measurements within the MRI scanner, adjustments were made to the previously mentioned in-vitro setup to align with the requirements of ferromagnetic-free equipment in the MRI scanner room (Fig. S6 and SI section S.6).

The gathered data underwent post-processing using MR analysis software Siswin64 (v0.95, Steffen Ringgaard, Aarhus University Hospital, Denmark), an in-house research software developed and available at the MRI research centre. Firstly, the MRI-integrated software provided both the geometrical and velocity information exported respectively as maps of pixel intensities and through-plane velocity components in the case of the 2D-flow. Then, in MATLAB, the velocity within the lumen were isolated through a threshold-based segmentation (Fig. S7 and SI section S.7) [16]. The velocity matrix was then considered for statistical analyses as described in the following section.

### 2.6 Statistical analysis of MRI and FSI results

To quantitatively assess the alignment of the digital twin outcomes with the MRI acquisitions, the FSI fluid velocity data were mapped onto the MRI structure grid through the FSI mesh spatial sampling, as done previously [29,30]. These transformations were performed in MATLAB, by extracting fluid velocities at the same spatial resolution as the MRI scans. Following this methodology, the FSI data were spatially downscaled from 0.65 mm to 1.1 mm. Fluid velocities were evaluated at three different cross-sections (Fig. 4), which were also acquired via MRI: one upstream of the valve (S1), one immediately downstream of the leaflets (S2), and one at the ring level of the model (S3). For each plane, **7** representative time instants were selected from the two datasets, starting from a sampling time of raw 2D-flow MRI data of 20 ms.

**Fig. 4.**
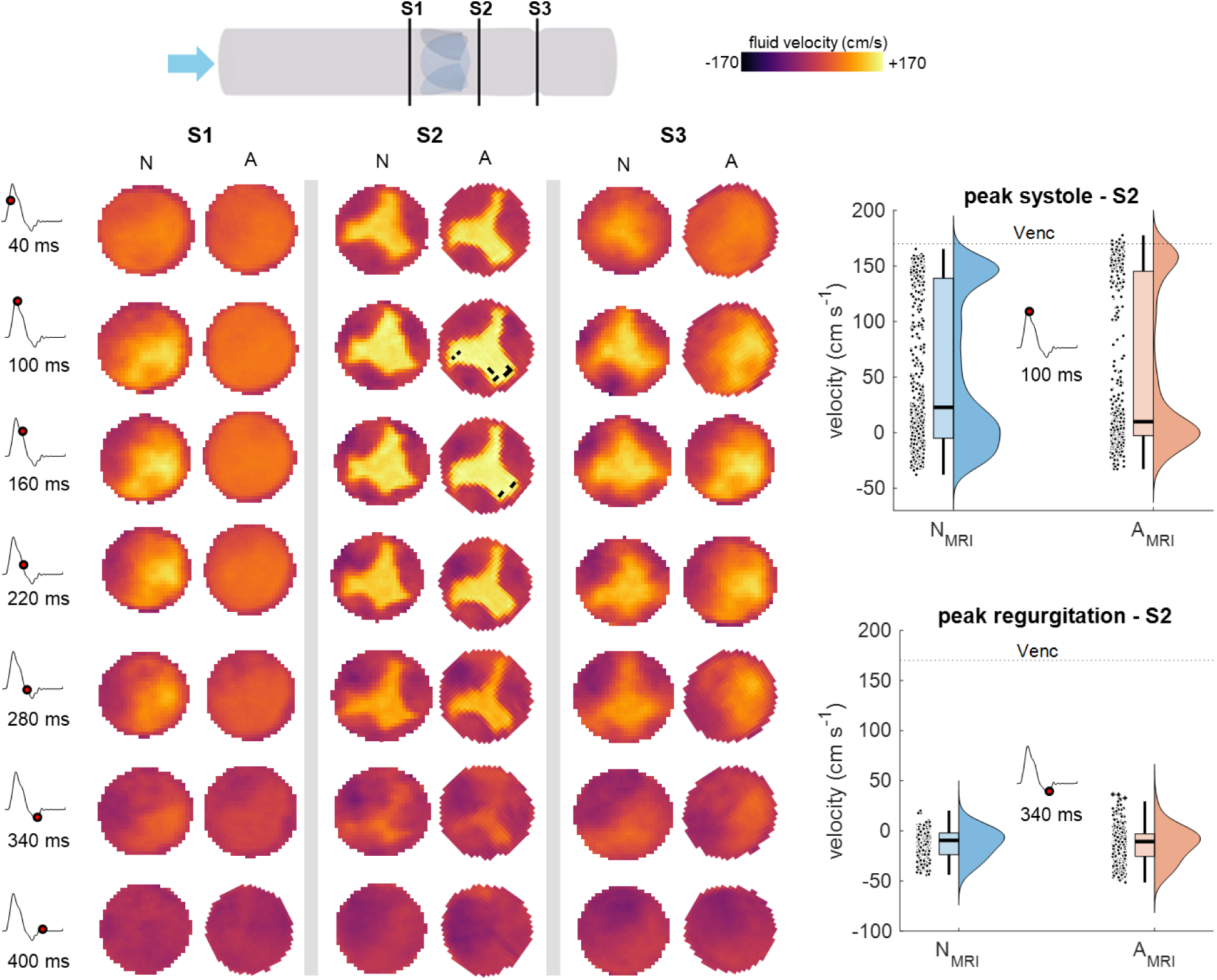
2D-flow MRI through-plane velocity contour plots for the native (N) and annuloplasty (A) 3D-printed phantom models. On the left, the time instant under consideration and on the top the representation of the location of the 3 cross-sections of interest. On the right, the distribution plots of cross-section S2 velocities of N and A models, considering the peak systole (t = 100 ms) and peak regurgitation (t = 340 ms) time points.

Both the MRI and FSI results were investigated in two fashions. First, the velocity distributions from both FSI and MRI data were compared. For individual cross-sections, differences between FSI and MRI were assessed with Kruskal-Wallis test and Mann-Whitney U-test, assuming significance for *p*-value < 0.05.

Next, given the large sample size (at least 500 data points), parametric methods were applied to analyze the velocity distributions. Mean values were extracted from these distributions and subsequently used for a global analysis, where a single representative mean velocity was determined for each cardiac time instant. Pearson’s correlation coefficient (R) was employed to measure the linearity between FSI and MRI velocities for both the native and annuloplasty models to provide a detailed comparison of the measured and predicted flow fields. Bland-Altman analysis was performed to investigate the agreement between FSI and 4D flow MRI data, with limits of agreements estimated as 1.45-fold the interquartile range (IQR). The average bias was estimated as the difference median. All the statistical analyses were performed in MATLAB.

## 3 Results

### 3.1 Effect of annuloplasty on aortic valve hemodynamics

The 3D-printed AR phantoms were successfully fabricated and connected to the flow-loop, through the connectors designed for the pressure readings. The flow rate curve depicted a full cardiac cycle, featuring a clear systolic peak at approximately 23.8±1.0 lpm (at t = 100 ms, Fig. S3 in SI). Additionally, the curve showed an average physiological flow rate of 5 lpm, with a regurgitation peak of 7.4±0.1 lpm (at t = 340 ms) before diastole. The 3D-printed valve of both the native and the annuloplasty models exhibited regurgitation. The pressure catheter positioned at the inlet of the phantom, upstream the valve, recorded a peak ventricular pressure of 118.2±3.3 mmHg during the systolic phase. Meanwhile, at the outlet, the aortic pressure ranged between a minimum of 78.0±0.5 mmHg and a maximum of 115.7±1.5 mmHg, resulting in normotensive systolic/diastolic aortic pressure readings of 115/78 mmHg. The difference between upstream and downstream valve pressures presented a maximum value of 2.5 mmHg during systole, as similarly reported [26].

Fig. 4 presents, on the left, the through-plane velocities quantified along the short-axis 2D-flow MRI acquisition for both the native and annuloplasty phantoms, considering the three cross-sections of interest. Given the chosen Venc of 170 cm/s, velocity values exceeding this threshold resulted in aliasing, as observed at t = 100 and t = 160 ms at cross-section S2 of the annuloplasty model [32]. From the qualitative viewpoint, the main differences between the models include a more rapid valve opening, as evidenced by the triangular jet shape emerging earlier (t = 40 ms) at cross-section S2 compared to the native model, and the occurrence of aliasing during peak systole (t = 100 ms) at S2 of the annuloplasty model. This initial analysis suggests that the maximum velocity reaches higher velocities than the Venc threshold during the systolic peak, possibly due to the area reduction imposed by the idealized implanted annuloplasty ring.

On the right, Fig. 4 shows the distribution plots at the systolic peak (t = 100 ms), chosen to avoid the geometry mismatch between the methods [29], and at the regurgitation peak (t = 340 ms). No statistically significant difference (*p*-value = 0.43) was observed between the velocity distributions of the native and annuloplasty models. However, at peak systole, the annuloplasty model exhibited a higher concentration of velocities near the 0-level (attributable to the presence of the ring) and around the Venc, whereas the native model showed a more dispersed velocity distribution between these two levels. During the regurgitation phase, the two distributions remained not statistically different (*p*-value = 0.11), though the annuloplasty model was characterized by a broader velocity distribution. The related median velocities and the IQR (75^th^ – 25^th^ percentiles) are provided in the SI section S.7. The respective flowrate values at peak systole and peak regurgitation, representing forward and backward flowrates, respectively, are considered more clinically relevant and are reported in Table 2.

**Table 2.** Forward and backward flowrate values for the native (N) and annuloplasty (A) MRI and FSI models, considered at peak systole (t = 100 ms) and peak regurgitation (t = 340 ms). Flowrates are shown as median and inter-quartile range (25^th^; 75^th^ percentiles).

|  | PEAK SYSTOLE<br>flow (lpm) |  | PEAK REGURGITATION<br>flow (lpm) |  |
| --- | --- | --- | --- | --- |
|  | N | A | N | A |
| **cross-section S1** |  |  |  |  |
| **MRI** | 20.9 (16.7; 23.5) | 19.4 (15.6; 22.3) | -4.1 (-5.6; -2.7) | -4.9 (-6.9; -3.0) |
| **FSI** | 20.4 (14.0; 24.5) | 21.0 (18.7; 22.8) | -4.1 (-15.1; 2.8) | -5.1 (-17.3; 2.7) |

| cross-section S2 |  |  |  |  |
| --- | --- | --- | --- | --- |
| **MRI** | 17.9 (-1.7; 47.7) | 17.4 (-0.9; 49.9) | -4.0 (-8.1; -0.7) | -4.5 (-8.8; -1.0) |
| **FSI** | 16.4 (0.0; 46.1) | 15.2 (-1.1; 43.9) | -12.1 (-16.4; -8.4) | -13.3 (-16.5; -10.0) |
| cross-section S3 |  |  |  |  |
| **MRI** | 13.9 (2.0; 27.1) | 15.1 (4.8; 26.0) | -3.3 (-7.5; 0.6) | -4.2 (-7.1; -1.3) |
| **FSI** | 13.3 (0.0; 31.8) | 18.0 (2.9; 32.7) | -11.5 (-16.9; -6.2) | -16.3 (-20.5; -12.2) |

### 3.2 Outcomes of the aortic annuloplasty digital twin

Before incorporating the experimental curves into the computational models as boundary conditions, the MCL acquisitions were filtered in MATLAB (Fig. 5), using a second-order notch filter at 50 Hz, and a third-order low-pass Butterworth filter with a cutoff frequency of 15 Hz to eliminate oscillations caused by the pump movement on the test bench. In terms of computational results, the simulations successfully captured the fluid flow within a conduit in the presence of a valve. To illustrate the evolution of fluid dynamics, the cross-sectional contour plots of the native and annuloplasty models at 7 time points during the cardiac cycle are presented in Fig. 5. Qualitatively, when analysing the post-annuloplasty model from the cross-sectional view, some alterations in the velocity patterns were observed compared to the native model. While upstream the valve (S1), the fluid velocities in the annuloplasty model do not qualitatively differ from the native model, alterations in velocity magnitudes are visible downstream (S2), becoming more evident at the ring level (S3), where velocity patterns are altered, as expected in the post-implantation conditions.

**Fig. 5.**
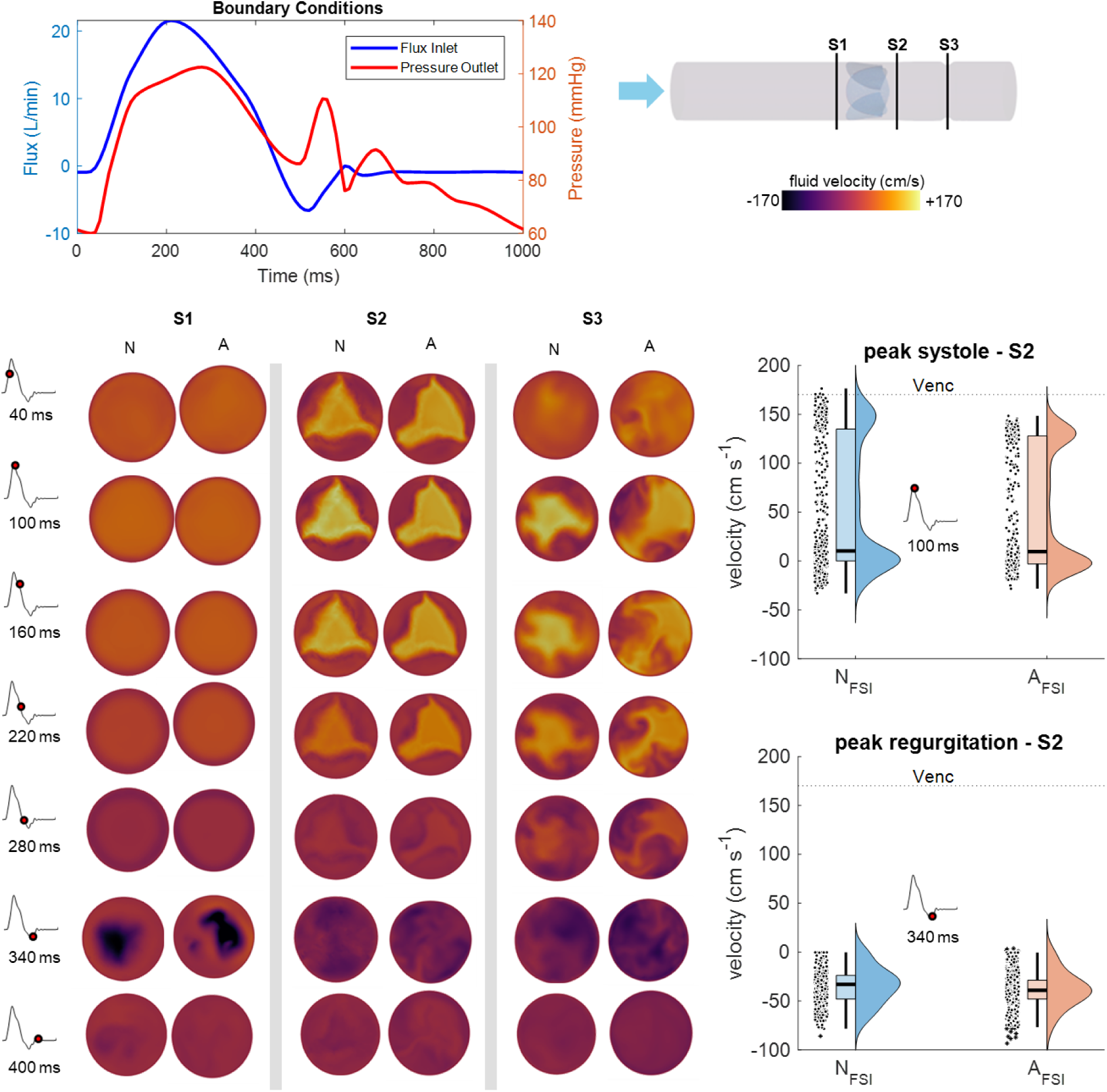
2D contour plots of the through-plane velocity components for the native (N) and annuloplasty (A) FSI models evaluated at the third cardiac cycle are reported. On the top, the inlet flow rate and outlet pressure curves, used as boundary conditions for the FSI simulations, and the representation of the location of the 3 cross-sections of interest are reported. On the right, the distribution plots of cross-section S2 velocities of N and A models, considering the peak systole (t = 100 ms) and peak regurgitation (t = 340 ms) time points.

The main differences were observed at the ring level (S3), where the presence of the annuloplasty ring led to alterations in the fluid flow, likely due to the diameter reduction, resulting in changes in the velocity magnitude patterns. This disruption became more pronounced after the systolic peak, with an increased formation of recirculation patterns of secondary flows becoming evident.

4D-flow MRI acquisitions of both the native and annuloplasty model showed the velocity intensities along with vectors (Fig. 6). With this imaging technique, the vectors display the magnitude of all three components of the fluid velocity. At peak systole (t = 100 ms), when the valve was fully open, the jet magnitude reached its maximum. At regurgitation peak (t = 340 ms), the backflow was clearly visible in both models, given the phantom insufficiency. However, the resolution of the scans did not allow the assessment of leaflets coaptation. Similarly to the native model, the opening and closing valve dynamics in the experimental setup remained consistent. In terms of qualitative assessment, non-negligible differences in the velocity patterns were observed between the native and annuloplasty MRI and FSI results. The presence of the annuloplasty ring, modelled as a diameter reduction, impacted both the jet flow during the systolic phase and the backflow during the regurgitation phase. Specifically, from the 4D-flow MRI image of the annuloplasty model at regurgitation peak, it is evident that the backflow was dampened.

**Fig. 6.**
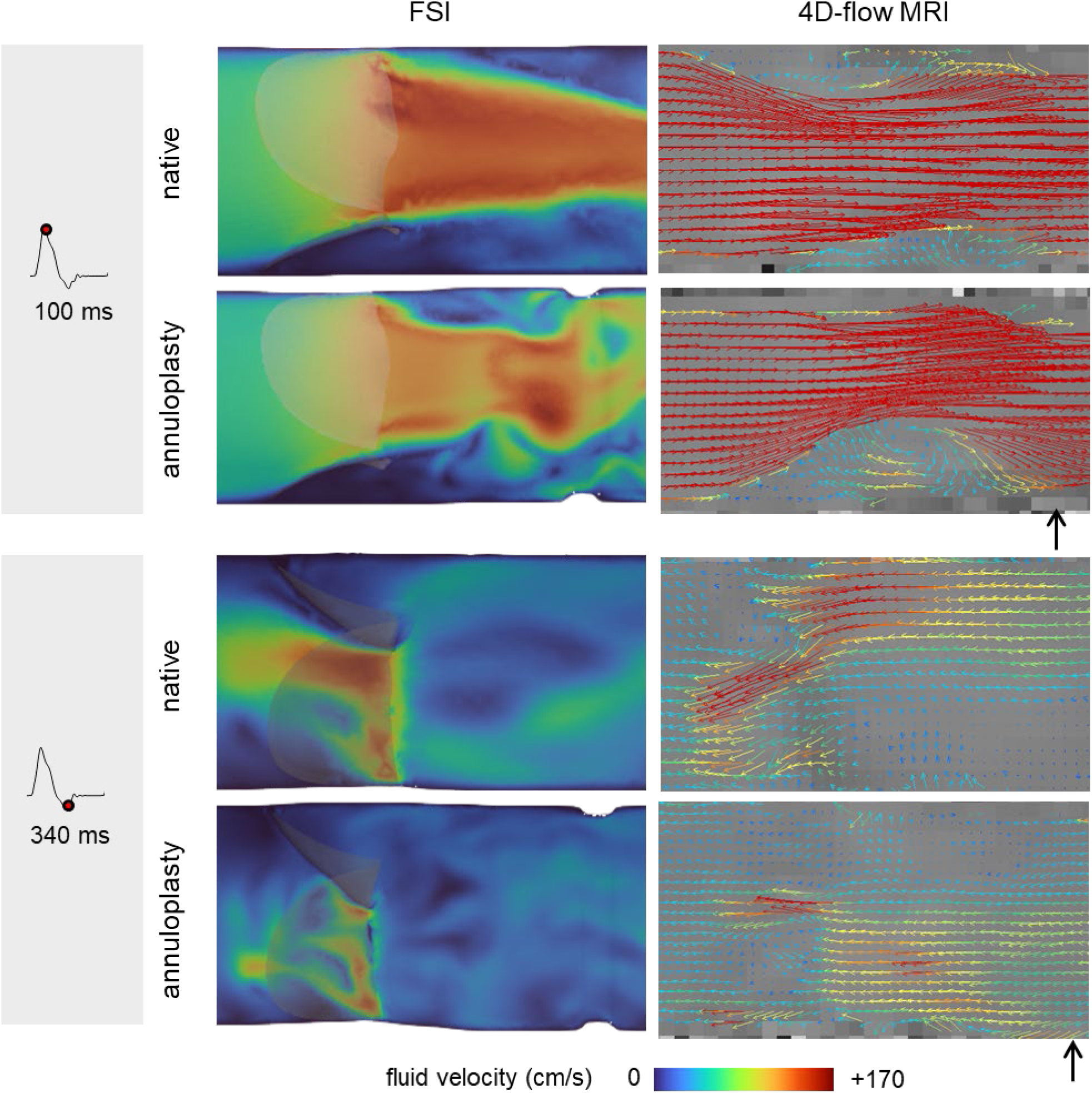
Qualitative comparison of FSI results and 4D-flow MRI velocity maps for both the native and annuloplasty models, evaluated at peak systole and peak regurgitation. In the FSI contour plots, the valve is shown for better understanding, while the black arrow indicates the location of the ring in the MRI vector maps.

### 3.3 Assessment of model accuracy

Overall, the FSI model successfully replicated the complex flow characteristics within the AR, showing qualitative agreement with 2D- and 4D-flow MRI measurements. A quantitative estimation of these findings is reported in Fig. 7, where the representative time-instant velocities per cross-section were extracted for both the MRI and FSI native and annuloplasty models. Despite the non-statistical significance, it is observable that (Fig. 7.A) the annuloplasty models showed higher velocities in forward flow phases, with exception of cross-section S1. On the contrary, during regurgitation and backward flow phases, the presence of the annuloplasty ring induced lower velocities as compared to the native scenarios. This finding suggests that the ring presence induces faster flows in both directions.

**Fig. 7.**
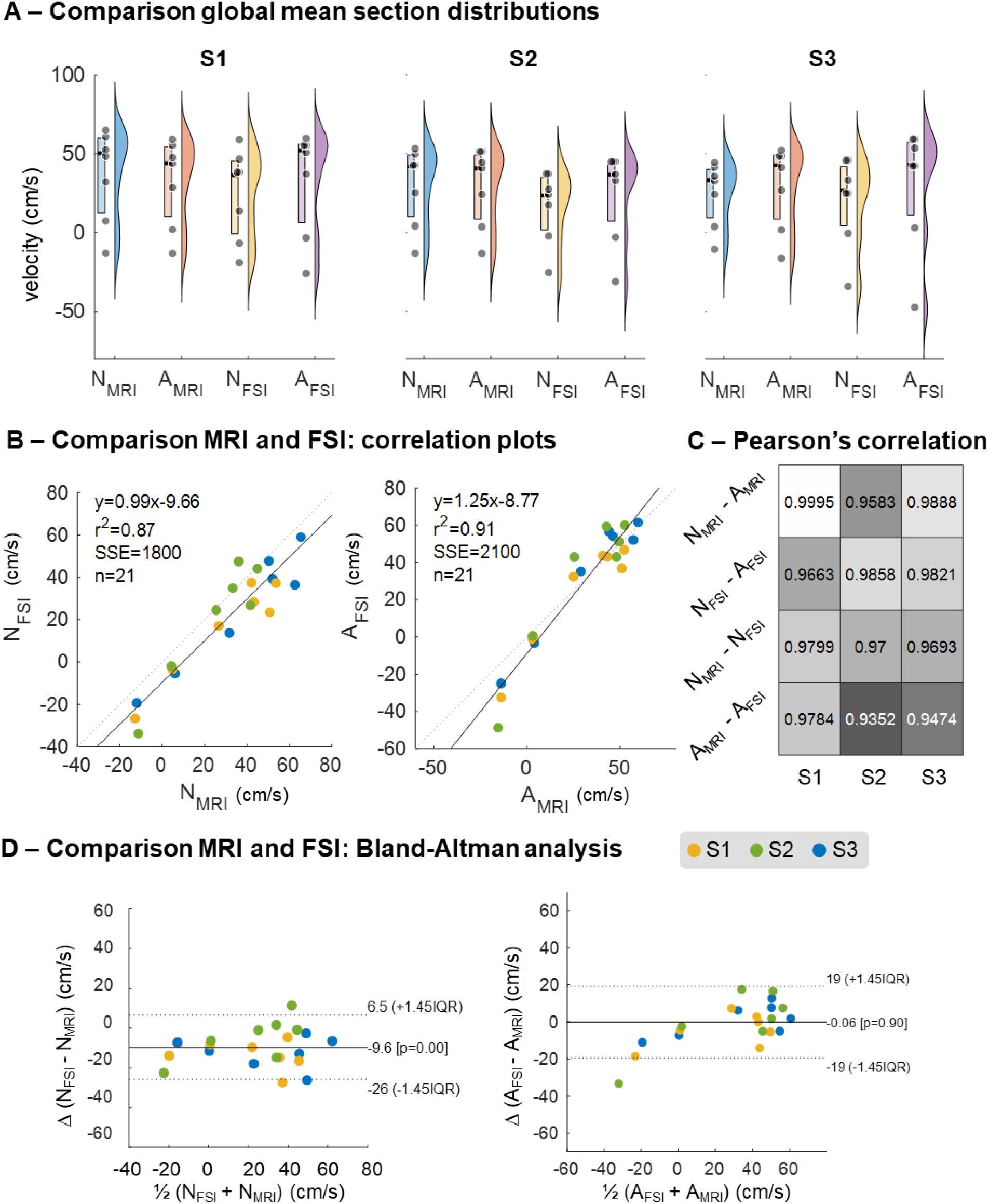
Results of the comparison of the native and annuloplasty model between the MRI and FSI velocities. Global values were considered (namely the mean velocity per section at the cardiac time instant). A) Distribution plots for the native and annuloplasty cases for the individual sections (the p-values of the significant differences are reported); B) Linear correlation plots for the native and annuloplasty MRI and FSI models for all the sections. C) Correlation map reporting the individual correlations for the global values. D) Non-parametric Bland-Altman comparison of the MRI and FSI native and annuloplasty models for all the sections.

The linear agreement between the methods, namely MRI and FSI, is demonstrated through correlation plots. Strong positive correlation, as globally shown in Fig. 7.B, was found for both native and annuloplasty models, with the correlation in the annuloplasty model being stronger than in the native model. The Pearson’s correlation map (Fig. 7.C) reveals that the lowest correlation (r = 0.9352) was found comparing the MRI and FSI annuloplasty model at cross-section S2. This lower correlation is attributed to the slight valve differences induced by the presence of the annuloplasty ring, as further supported by the individual cross-section results reported in SI section S.7 and shown in Fig. S7.1-3.

Finally, as reported in Fig. 7.D, Bland-Altman analyses proved that the difference between the annuloplasty MRI and FSI models was global null, while the MRI provided larger velocities for the native model. Regarding the annuloplasty model, the largest difference was observed during the regurgitation phase, with the FSI model exhibiting significantly higher regurgitation.

## 4 Discussion

This study evaluated the hemodynamic impact of aortic annuloplasty (AA) using an integrative methodology combining 3D-printed phantom models, in-vitro flow-loop experiments, computational simulations, and advanced imaging techniques. The findings highlight the ability of this approach to provide both qualitative and quantitative insights into flow alterations induced by AA, contributing to a better understanding of its implications for AV function.

The 3D-printed phantoms successfully replicated the native and annuloplasty conditions [33,34], allowing for controlled experimental measurements in the MCL. The observed hemodynamic parameters were consistent with physiological values, with peak ventricular pressures of approximately 117.8 ± 3.1 mmHg and systolic/diastolic aortic pressures of 115/78 mmHg. These results validate the experimental setup as a representative model of native and post-AA conditions [7,9,10].

The annuloplasty model introduced notable changes in flow dynamics. Specifically, the reduced diameter imposed by the annuloplasty ring was associated with higher peak systolic velocities, evidenced by the triangular jet pattern observed earlier in the cardiac cycle compared to the native model. The aliasing seen during peak systole (e.g., at cross-section S2) underscores the velocity magnitude exceeding the encoding threshold, suggesting localized acceleration due to the ring’s geometrical constraints. Interestingly, despite these differences, no statistically significant variations in velocity distributions were detected between the native and annuloplasty models at peak systole or regurgitation. However, a broader velocity distribution during regurgitation was observed in the annuloplasty model, potentially reflecting increased complexity in backflow patterns.

The FSI-based digital twin successfully replicated the velocity distributions observed experimentally. The velocity patterns at cross-section S3 demonstrated significant alterations attributable to the annuloplasty ring, with recirculation zones and secondary flow structures becoming prominent downstream of the ring during late systole. These features were less pronounced in the native model, highlighting the influence of ring-induced geometric constraints. Correlation analysis between FSI and 2D- and 4D-flow MRI measurements revealed strong agreement, with Pearson correlation coefficients exceeding 0.93 for all configurations. Notably, the weakest correlation was found at cross-section S2 of the annuloplasty model, likely due to discrepancies arising from the valve’s altered geometry and ring-induced flow acceleration. Bland-Altman analysis further confirmed the consistency between methods, although the annuloplasty model exhibited higher regurgitant velocities in the FSI simulations compared to MRI measurements. This discrepancy may stem from limitations in experimental imaging resolution [35], particularly during regurgitation phases.

The observed increase in peak systolic velocities and altered recirculation patterns in the annuloplasty model underline the potential hemodynamic consequences of AA. While these changes may not directly affect immediate valve performance, they could have implications for long-term AV durability or the development of secondary flow-related complications. The ability of the digital twin to predict these changes highlights its potential as a tool for preoperative planning and postoperative assessment, offering clinicians a non-invasive tool to optimize AA designs and patient-specific treatment strategies.

This was the first known integrative study on AA. Therefore, many challenges were faced during the process of combining experimental data, simulations analysis, and clinical imaging.

Regarding the experimental work, assumptions in the designing and manufacturing process of the phantoms were made. Enhancing the design of the AR to incorporate more anatomical features, such as the inclusion of the Valsalva sinuses’ structure, could offer a more accurate representation of its dynamics, particularly in relation to the AV. However, this may pose challenges in the 3D- printing process. On the computational front, simulations need refinement to better replicate the experimental observations and imaging data. Specifically, additional numerical tuning of alternative leaflet contact algorithms and contact parameters could improve the simulation results. This could enable quantitative measurements of leaflet coaptation from the computational model [38,39], crucial for assessing AA experimental conditions. Furthermore, conducting simulations with the developed AA model could assess various positions of annuloplasty ring implantation. This would enable clinicians to evaluate annuloplasty outcomes more effectively, as assessing the positioning of the ring is challenging experimentally.

The discrepancies in velocity results between the FSI simulations and 4D-flow MRI data may arise from two main issues, both related to the 4D-MRI scans’ relatively low image quality [35]. Firstly, due to the scan parameters – specifically, the Venc settings, which control the range of velocities accurately captured, and thereby velocity sensitivity – some high or low velocities might not be fully represented. To address this issue, a dual Venc approach could be employed, where two separate scans capture a broader range of velocities. However, this method is often impractical in clinical environments because it significantly increases the required scanning time [40]. Secondly, during the resolution downscaling process of FSI results, data sampling could introduce a numerical bias. The sampling rate should be then evaluated through an impact assessment on the fluid velocities resulting from the downscaling.

## 5 Conclusions

The integrated framework presented in this work allowed comparing AA from the hemodynamic viewpoint, through the virtual implantation of the ring in an idealized AR digital twin and through 4D-flow MRI scans in an idealized AR 3D-printed phantom.

Firstly, significant progress was made in the experimental analysis of AA. This included the development of an idealized model of native AR and AA, optimization of the 3D-printing process for compliant AR fabrication, and integration with the flow-loop. Furthermore, the acquisition of sensor-based flow rate and pressure data for the outlined experiments ensured robust data collection and thorough analysis. Secondly, the study conducted the first known FSI analysis of idealized post-annuloplasty conditions of the AR, providing proof of concept for the methodology, while acknowledging strong assumptions in geometry and boundary conditions, and recognizing that the simplified models currently applied have too many limitations to draw general conclusions about the impact of AA procedures on in vivo hemodynamics. Thirdly, the experimental setup was optimized for MRI scans, allowing for comprehensive analysis of 2D-flow and 4D-flow MRI data. The flow velocity distributions were well correlated between FSI and MRI outcomes, as shown by Pearson’s correlations and Bland-Altman analyses.

By combining the flexibility and high-resolution capabilities of computational modelling with rigorous validation procedures, computational simulations play a key role in advancing the understanding of AA’s impact on AR hemodynamics and informing clinical decision-making processes. This study serves as an important stepping stone toward achieving these goals.

## Acknowledgements

This work acknowledges Johannes Høgfeldt Jedrzejczyk, Rahima Khouani and Stefine Varde for the help in performing experiments at Aarhus University Hospital.

## Author contributions

**LB**: Conceptualization, Methodology, Formal analysis, Investigation, Data curation, Writing – Original Draft, Writing – Review and Editing, Visualization. **MZ**: Formal analysis, Investigation, Data curation, Writing – Review and Editing. **AR**: Formal analysis, Investigation, Data curation, Writing – Review and Editing. **FM**: Supervision, Writing – Review and Editing. **SR**: Conceptualization, Investigation, Data curation, Writing – Review and Editing. **WYK**: Conceptualization, Writing – Review and Editing, Funding acquisition. **PJ**: Conceptualization, Supervision, Methodology, Project administration, Writing – Review and Editing, Funding acquisition. **MC**: Conceptualization, Methodology, Formal analysis, Investigation, Data curation, Writing – Original Draft, Review and Editing, Supervision, Project administration, Funding acquisition.

All authors contributed to the article and approved the submitted version.

## Acronyms

AV: aortic valve
AR: aortic root
AA: aortic annuloplasty
FSI: fluid-structure interaction
MRI: magnetic resonance imaging
CAD: computer aided design
MCL: mock circulatory flow-loop

