## Supplementary Materials for "Aortic annuloplasty FSI digital twin of 3D-printed phantoms with 4D-flow MRI comparison"

### S.1 Fabrication of 3D-printed AR phantoms

The AR phantoms were created using resin-based stereolithography 3D printing, which employs photopolymerization to build objects layer by layer. CAD models were designed in SolidWorks 2022 (Dassault Systèmes SolidWorks Corp., Waltham, MA, USA) and exported as .STL files to PreForm (Formlabs Inc., Somerville, MA, USA) for support optimization and orientation. A flexible resin, Elastic 50A (Formlabs Inc., Somerville, MA, USA) was employed in the Form 3B+ printer (Fig. S1A) in the MAKERLab Pro facility (Aarhus University, Department of Electrical and Computer Engineering). Following printing, samples were rinsed in isopropyl alcohol for 20 minutes and UV-cured at 60°C for 20 minutes, adhering to the manufacturer's postprocessing instructions.

Achieving the physiological leaflet thickness of 0.1–0.2 mm [1] was deemed unfeasible with the available equipment after several printing trials. To address this, an iterative approach was employed, starting with a leaflet thickness of 0.1 mm and increasing by 0.05 mm in each attempt. Leaflets up to 0.2 mm thickness failed to form properly. At 0.35 mm, the leaflets were successfully printed but presented challenges in removing support material due to the elastic behaviour of the resin before UV-curing. Ultimately, a thickness of 0.4 mm was adopted, allowing the successful fabrication of multiple AR phantoms while ensuring both structural integrity and ease of postprocessing.

Similarly, producing the phantom in a single printing process proved impractical due to the difficulty of removing support material inside the cylindrical conduit, which was designed to stabilize the valve. To address this, the upstream and downstream sections of the cylinder were printed separately from the valve (Fig. S1B). Before post-curing, the three components were assembled using Elastic 50A resin as an adhesive. The final assembly underwent UV-light treatment to ensure robust bonding, without altering mechanical properties of the resin.

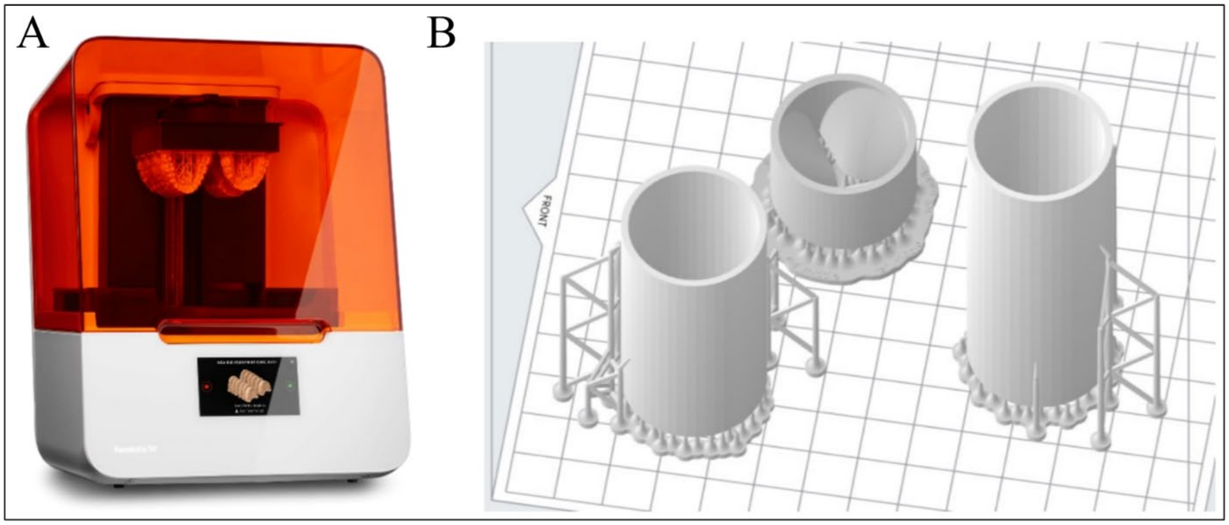


**Fig. S1** – (A) Form 3B+ printer (Formlabs Inc., Somerville, MA, USA); (B) PreForm view of the upstream, valve, downstream sections with the support material.

### S.2 50A Resin mechanical characterization

Uniaxial tensile testing of the Elastic 50A Resin (Formlabs Inc., Somerville, MA, USA) was performed at MaterialsLab facility (Aarhus University, Department of Mechanical and Production Engineering) in order to evaluate the Young's modulus of the material. Tests were conducted according to ISO 37-1:2017 guidelines for the determination of tensile stress-strain properties of rubber, vulcanized or thermoplastic materials. Three dumbbell test pieces (Figure S2A) were stretched in a 50 kN Zwick tensile testing machine (Z050, ZwickRoell GmbH & Co. KG, Ulm, Germany) at a constant rate of 100 mm/min of traverse of the driven grip.

The steps of the testing protocol are reported below.

- *Measurement of the thickness at the centre and at both ends of the dumbbell.*
- *Positioning of the piece into the tensile-testing machine.*

- *Verification that the end tabs are gripped symmetrically to ensure that the tension is distributed uniformly over the cross-section.*
- *Application of a 0.1 MPa prestress to avoid bending of the dumbbell when the initial test length is measured.*
- *Setting of the nominal rate of traverse for the moving grip at 100 mm/min.*
- *Start of the machine.*
- *Monitoring of the change in length of the dumbbell and force throughout the test until breakage of the piece.*
- *Repetition for three test pieces.*

The experimental data collected was post-processed in MATLAB. The results of the testing of the three test pieces are reported in Figure S2B. Stress and strain data were obtained from the experimental data by considering the width, thickness and initial length of the dumbbells (Figure S2C). The experiments confirmed the tensile non-linear behaviour of the Elastic 50A resin. However, an initial linear region for small strains below 0.25 could be identified and it was used to calculate the Young's modulus by means of a linear fitting [2]. The Young's modulus resulted of 1.99 MPa, which is coherent with the mechanical properties of Elastic 50A Resin reported from the manufacturer [3].

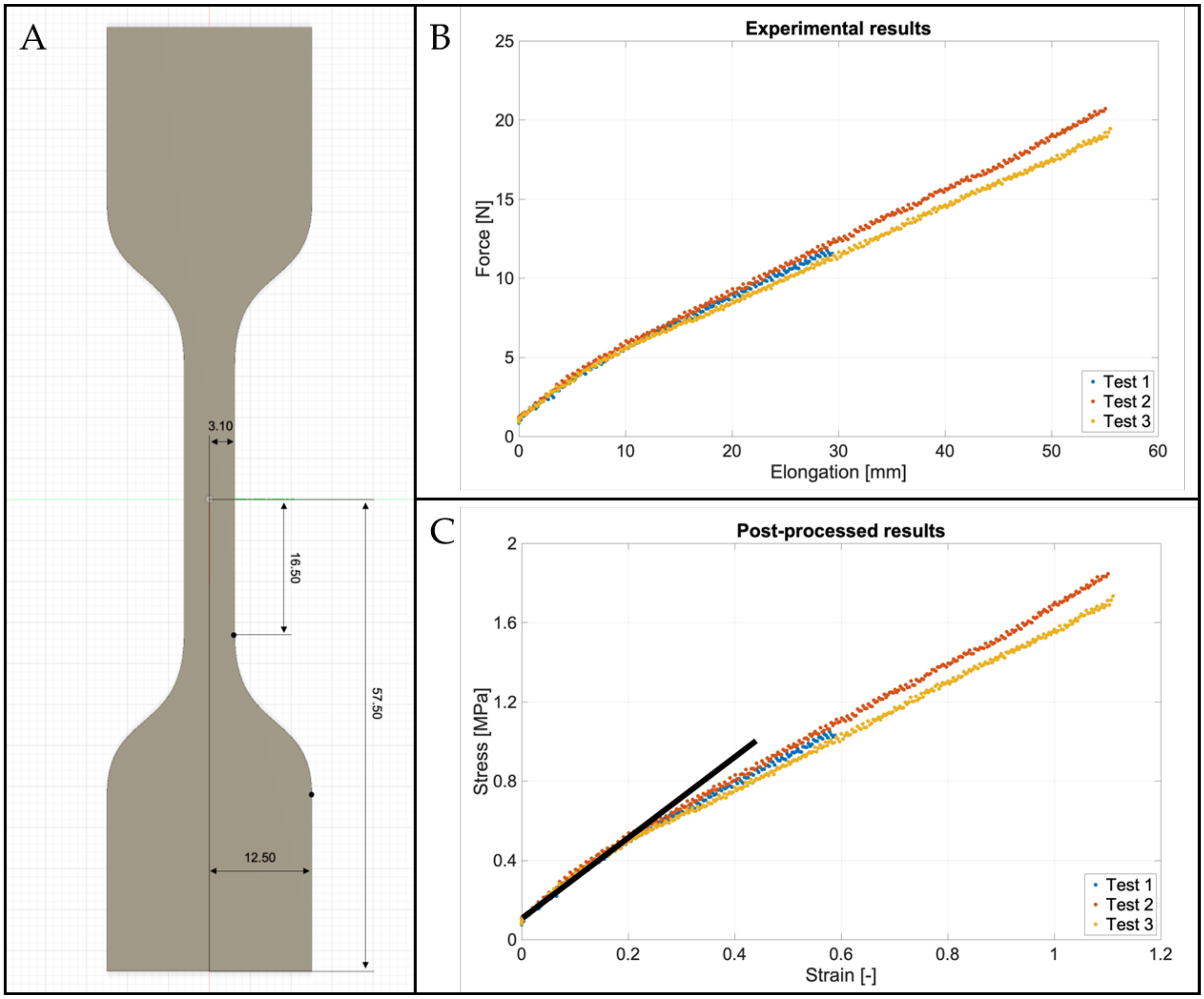


**Fig. S2** – (A) Final geometry of the test piece with measurements in mm; (B) force-displacement behaviour of the three dumbbells during tensile testing; (C) stress-strain behaviour of the three dumbbells during tensile testing.

### S.3 Sensors-based collected data processing

To ensure statistical reliability, a series of experiments were carried out for both the native and annuloplasty models. For each model, three independent tests were conducted on separate days. In each test session, the model, sensors, and pump were connected to the MCL. Within each test, three consecutive acquisitions were performed. Each acquisition involved the recording of five cardiac cycles.

#### S.3.1 Example of raw data

In this section, an example of the raw data collected during the experiments for both phantoms is reported (Figure S3). Data is presented as mean $\pm$ standard deviation.

The flowrate curves display a complete cardiac cycle lasting 1000 ms, characterized by a distinct systolic peak around 20.1$\pm$0.3 lpm and 23.8±1.0 lpm (at t = 100 ms), for the native and the annuloplasty phantom, respectively. The curves also exhibit a mean physiological flowrate of 5 lpm (corresponding to a stroke volume of the pump of 83.3 ml/s), with regurgitation exceeding 5 lpm prior to diastole (at t = 340 ms). The obtained regurgitant flowrate of 6.2$\pm$0.2 lpm and 7.4±0.1 lpm, respectively was not comparable to physiological values, meaning that both the 3D-printed valves were insufficient.

The ventricular pressure measured using a fluid-filled pressure catheter positioned at the inlet of the phantoms, upstream the valve, exhibited a peak of 116.0$\pm$4.5 mmHg and 117.8±3.1 mmHg, respectively, during the systolic phase. Throughout the diastole, the pressure remained relatively constant around the zero value, though the graph was shifted by approximately 10 mmHg due to the zero-pressure condition of the setup (pressure drop of the water column present in the ventricular and aortic chambers).

The averaged aortic pressure, measured at the outlet of the phantoms, ranged between a minimum of 69.8$\pm$2.5 mmHg and a maximum of 118.2$\pm$3.3 mmHg for the native phantom, and between a minimum of 78.0$\pm$0.5 mmHg and a maximum of 115.7$\pm$1.5 mmHg for the annuloplasty phantom, consistently with a physiological aortic pressure during the whole cardiac cycle. Normotensive conditions, characterized by systolic/diastolic aortic pressure readings of 118/69 mmHg and 115/78 mmHg, respectively, were obtained through the fine-tuning of mechanical resistance on the venous return tube and the regulation of air volume within the compliance chamber.

Furthermore, it is worth noting that the experimental conditions replicated in terms of flow rate and pressure with the MCL in CAVE Lab were consistent with *in vitro* testing operated with different MCLs [4–7].

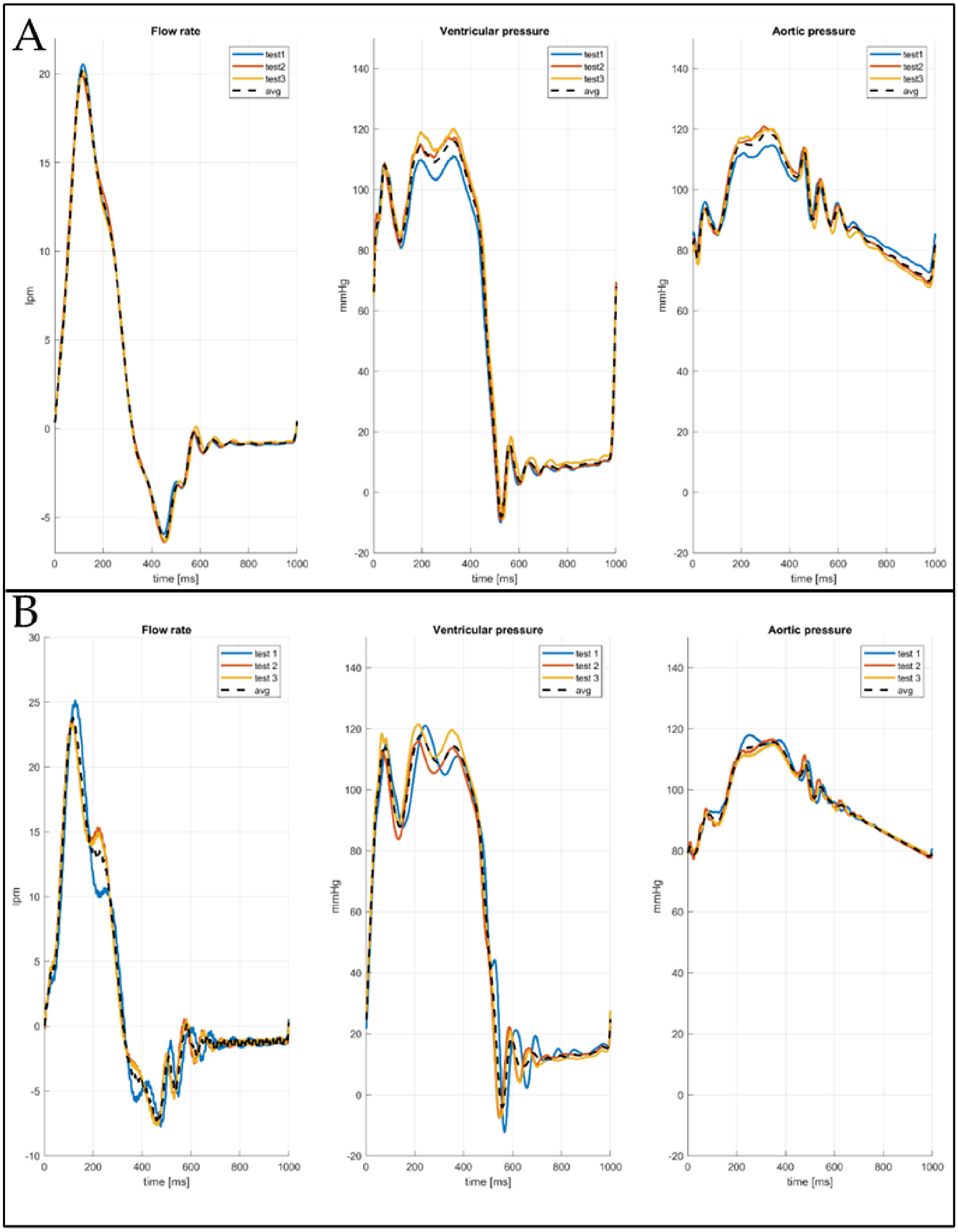


**Fig. S3** – Experimental curves for the native model (A) and the annuloplasty model (B): sensors-based collected data of flowrate, ventricular and aortic pressures obtained through averaging of the tests.

#### S.3.2 Data post-processing to create boundary conditions

Following the data collection using LabVIEW, additional processing steps were undertaken to refine the data before its utilization in the computational model. This involved implementing noise filtering, in particular, the removal of 50 Hz noise by a second order lowpass Butterworth filter, and a third-order low-pass Butterworth filter with a cutoff frequency of 15 Hz to eliminate oscillations caused by the pump movement on the test bench. To create a file to use as boundary condition curve in the computational model, the flow rate data underwent processing to extract the corresponding velocity magnitude curve. Following this, the velocity and pressure values were converted to I.S. units to ensure conformity with the unit system adopted in the simulation software. The averaged velocities and pressures, derived as mean values over cardiac cycles, acquisitions, and tests, were systematically repeated three times to construct a comprehensive file representing three complete cardiac cycles.

### S.4 Mesh sensitivity analysis

Four different shell meshes, ranging from coarsest (M0) to finest (M3), were generated to conduct a sensitivity analysis on the discretization density. Since the fluid volume discretization is generated automatically by LS-Dyna (Ansys, Canonsburg, PA, USA), starting from the structural mesh, the mesh sensitivity analysis is performed on the discretization of the valve and the cylindrical structure. Each mesh differed from the previous one by approximately 65$\pm$2% in the number of elements. In Figure S4.1A, the four meshed AVs are presented to illustrate the increase in the number of elements, resulting in a decrease in element size with each meshing iteration.

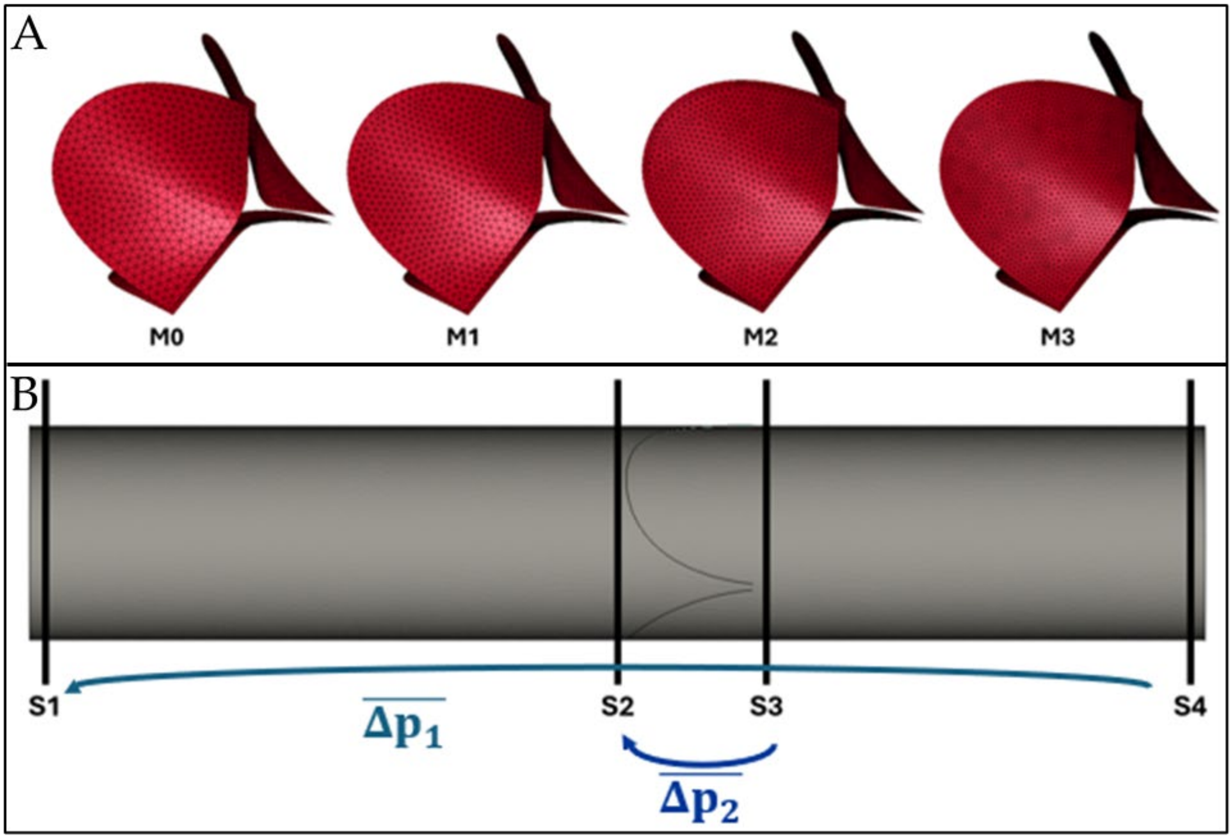


**Fig. S4.1** – **A**) Leaflets meshes for mesh sensitivity analysis; **B**) Cut planes chosen for mesh sensitivity analysis, average pressure drop across the whole model ($\overline{\Delta p_1}$) and average transvalvular pressure drop ($\overline{\Delta p_2}$).

Given that boundary conditions do not directly impact the accuracy of mesh sensitivity, simulations conducted for this purpose were executed with simplified boundary conditions to mitigate computational costs. A literature velocity profile was imposed at the fluid inlet [8], while a constant zero pressure condition was chosen for the outlet, as done previously [9,10].

To monitor the convergence of the mesh, four sections of the fluid volume along the longitudinal direction of the cylindrical structure were chosen (Figure S4.1B). These cut planes were positioned near the inlet (S1) and outlet (S4) of the fluid volume, as well as upstream (S2) and downstream (S3) the AV. Average velocity and average pressure in these cut planes were assessed throughout the entire cardiac cycle to evaluate the convergence of results.

When a mesh size of 0.65 mm was set (M2), the mismatch with respect to the results obtained with a 0.5 mm mesh size (M3) fell below an acceptable threshold. The comparison of average velocity (Figure S4.2A), average pressure drop across the whole model and average transvalvular pressure drop (Figure S4.2B) between the different geometry discretization was evaluated using a percentage error, defined as:

$$err\% = \frac{|x_{i-1} - x_i|}{|x_{i-1}|} \times 100 < 5\%$$

Where the results $x$ from the i-th mesh are considered. Based on these results, a mesh with a 0.65-mm element size was considered adequately refined to guarantee mesh independency of computed results and was set for each AR model.

In order to balance the computational efficiency and capturing essential dynamics, three cardiac cycles were simulated. This strategic choice arises from the observation that the fourth cycle introduced minimal variation (Figure S4.3A), making the inclusion of additional cycles computationally expensive without significantly enhancing the insights gained from the simulation results.

Convergence of the mentioned parameters was achieved with the second finer mesh, i.e., M2. This mesh was subsequently chosen as the geometry discretization for both native and annuloplasty models, as further mesh refinement was deemed to have a non-significant impact on results. The total computation time for each simulation, conducted on 40 CPUs of an Intel Xeon64 system with 240 GB of RAM, was also a factor under consideration. Notably, as depicted in Figure S4.3B, M2 required approximately 45% less computation time compared to the finer mesh, M3.

A summary of the geometry discretization details for the native and annuloplasty AR models is reported in Table S1. The table lists the total number of structural and fluid elements used in each model. The structural mesh includes 3-node shell elements, while the fluid volume is represented by tetrahedral Eulerian elements.

**Table S1** – Number of elements used to spatially discretise the structural part (AV, AR, inlet and outlet) and the fluid volume for the native (N) and annuloplasty (A) models.

**Spatial discretization**

| | Structural mesh (3-node shell) | | | | | Fluid volume mesh (tetrahedral) |
|---|---|---|---|---|---|---|
| | AV | AR | INLET | OUTLET | TOT | FLUID |
| **N** | 5,740 | 65,607 | 2,941 | 2,884 | 77,172 | 1,9M |
| **A** | 5,740 | 67,592 | 2,941 | 2,884 | 79,361 | 2,05M |

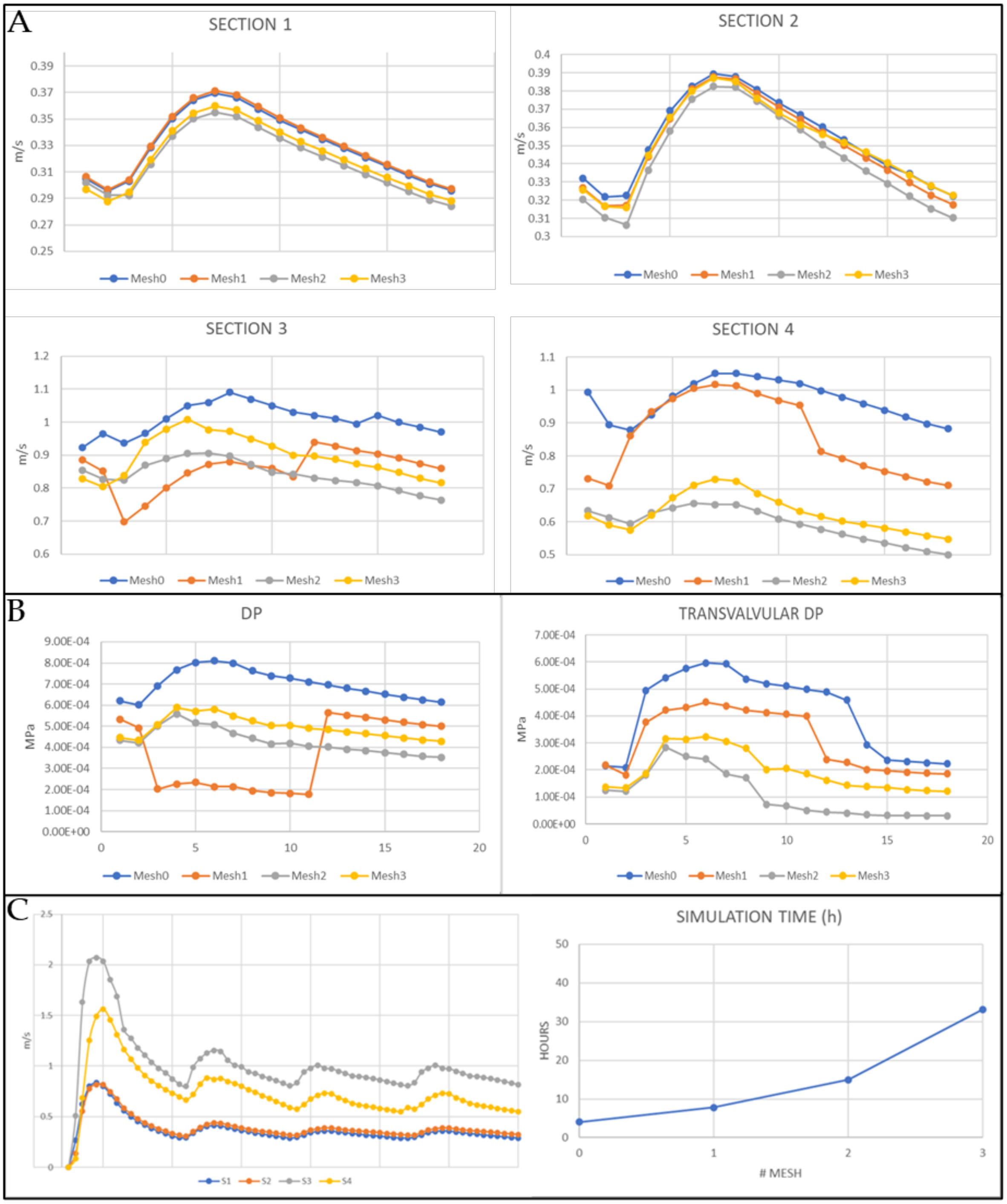


**Fig. S4.2** – **A**) – Average velocity obtained at different cut planes with the four different meshes; **B**) average pressure drop across the whole model and average transvalvular pressure drop obtained at different cut planes with the four different meshes; **C**) average velocity obtained at different cut planes with M2 during four cardiac cycles and simulation time with the four different meshes.

### S.5 MRI acquisition setup

To facilitate measurements within the MRI coil, adjustments were made to the previously mentioned in vitro setup to align with the requirements of ferromagnetic-free equipment in the MRI scanner room. These modifications involved using longer tubes (~10 m), incorporating sliding valves, and employing pressure data collection via long fluid-filled catheters (~10 m) to place all the metallic components in the controller room through ad hoc openings in the wall and allow for remote control. To synchronize the piston pump with the MRI sequences, a built-in trigger signal from the pump was used as a wireless ECG signal for the MRI machine.

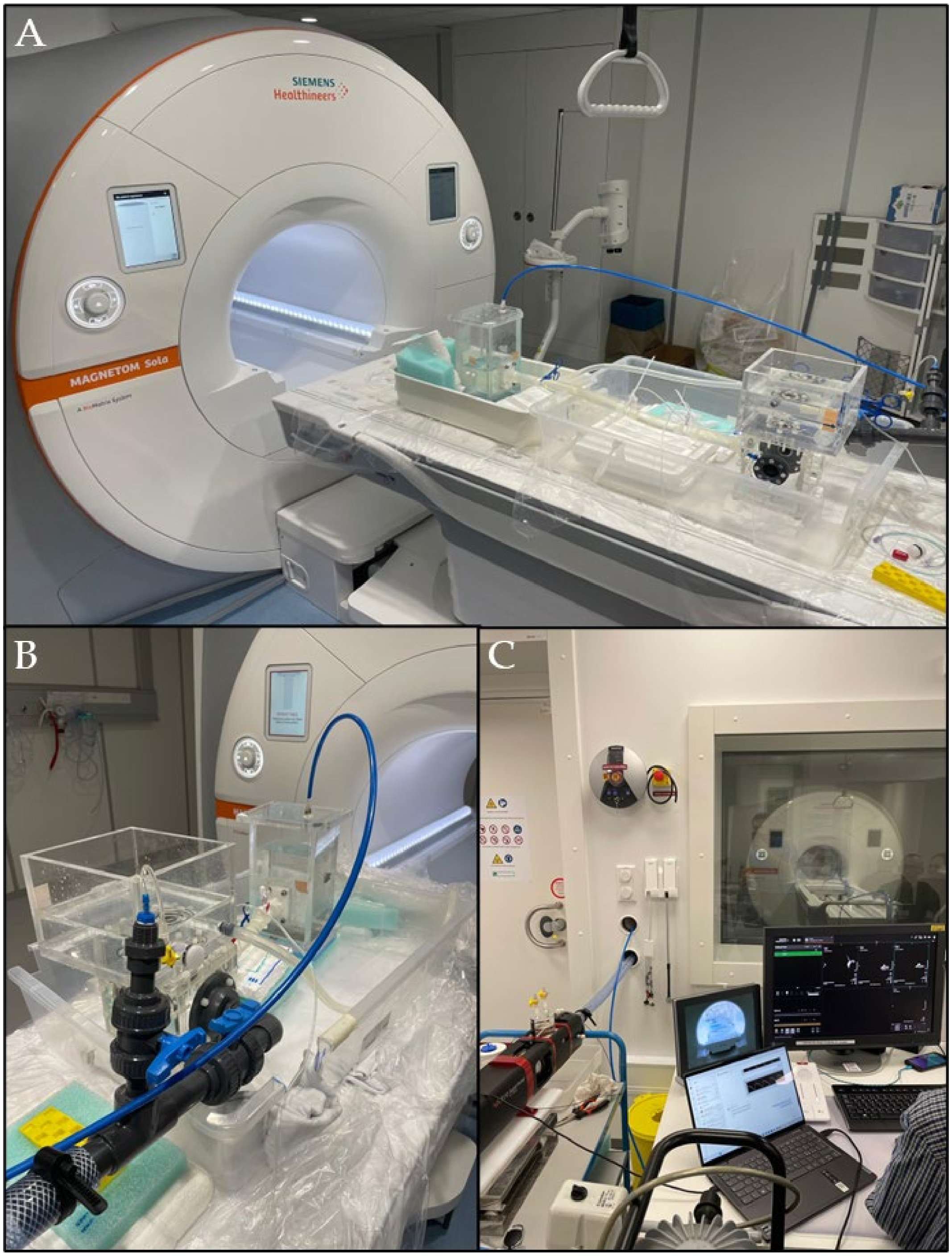


**Fig. S5** – **A)** CAVE Lab MCL in the MRI scanner at Aarhus University Hospital; **B**) air vents and valve system implemented to adapt MCL for MRI-compatibility; **C**) remote control of MCL through MRI control room.

### S.6 Post-processing of MRI scans

The MRI-integrated software provided images solely containing geometrical information, necessitating post-processing with separate software to extract fluid velocity magnitude distribution. Employing an available in-house MATLAB code [11], grey-scale fluid velocity maps were subsequently coloured according to the selected colour scale (Fig. S4). Given the chosen enclosure velocity of 1.7 m/s, the colour scale used in the visualization was adjusted accordingly, with the maximum colour intensity corresponding to this velocity value. When the velocity value in the MRI images exceeded the set threshold, aliasing phenomenon occurred [12].

To remove the noise outside the lumen (shown as the random coloured pixel), the binary image of the geometrical information (Fig. S6 on the left) was used to select the inside-lumen pixels. This information was collected in an array of points later used for quantitative considerations.

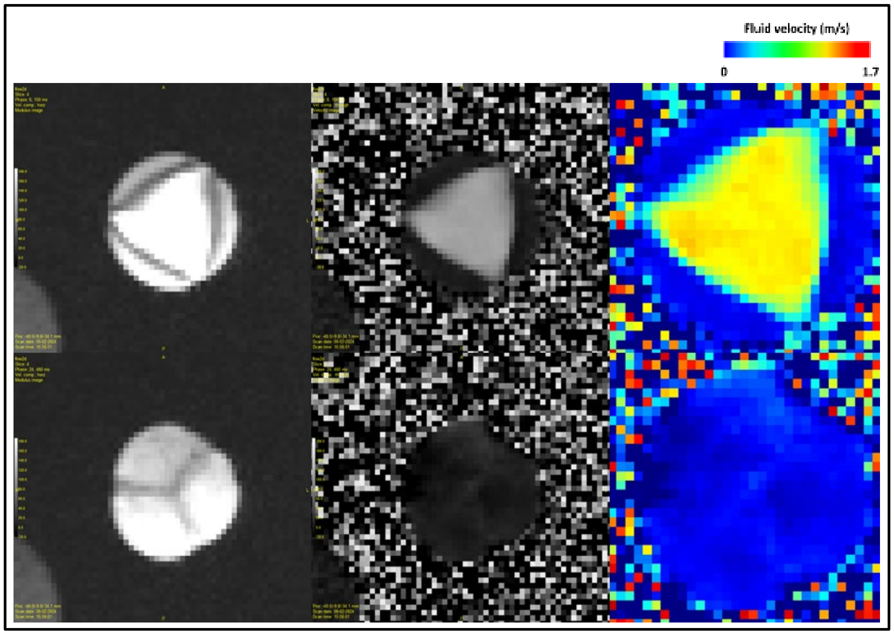


**Fig. S6** – Postprocessing of the MRI scans: on the left, original geometry images from the 2D-flow scans; in the middle, original fluid velocity grey-scale intensity profile; on the right, processed and coloured fluid velocity contour plot.

### S.7 Statistical results of MRI-FSI agreement

The median velocities and the IQR (75$^{th}$ – 25$^{th}$ percentiles) for both native and annuloplasty MRI and FSI data at peak systole and peak regurgitation are reported in Table S2.

**Table S2** – Velocities values for the native (N) and annuloplasty (A) MRI and FSI models, considered at peak systole (t = 100 ms) and peak regurgitation (t = 340 ms). Velocities are shown as median and inter-quartile range (25$^{th}$; 75$^{th}$ percentiles).

| | **PEAK SYSTOLE velocity (cm s$^{-1}$)** | | **PEAK REGURGITATION velocity (cm s$^{-1}$)** | |
|---|---|---|---|---|
| | **N** | **A** | **N** | **A** |
| **cross-section S1** | | | | |
| **MRI** | 65.5 (48.6; 68.5) | 59.7 (45.4; 65.1) | -12.0 (-16.5; -8.1) | -13.9 (-20.1; -8.9) |
| **FSI** | 66.5 (41.0; 71.4) | 62.8 (54.5; 66.4) | -0.0 (-44.2; 8.4) | -1.9 (-50.6; 8.0) |
| **cross-section S2** | | | | |
| **MRI** | 22.7 (-5.1; 139.0) | 9.8 (-2.7; 145.4) | -9.5 (-23.8; -2.2) | -10.9 (-25.7; -2.9) |
| **FSI** | 9.2 (0.0; 134.2) | 8.4 (-3.2; 128.0) | -33.3 (-48.0; -24.6) | -39.3 (-48.3; -29.3) |
| **cross-section S3** | | | | |
| **MRI** | 45.4 (7.5; 98.9) | 52.5 (17.7; 94.6) | -11.6 (-27.6; 2.2) | -15.5 (-25.9; -4.9) |
| **FSI** | 30.0 (0.0; 115.7) | 67.5 (10.8; 119.0) | -41.8 (-61.7; -22.6) | -59.1 (-74.6; -44.7) |

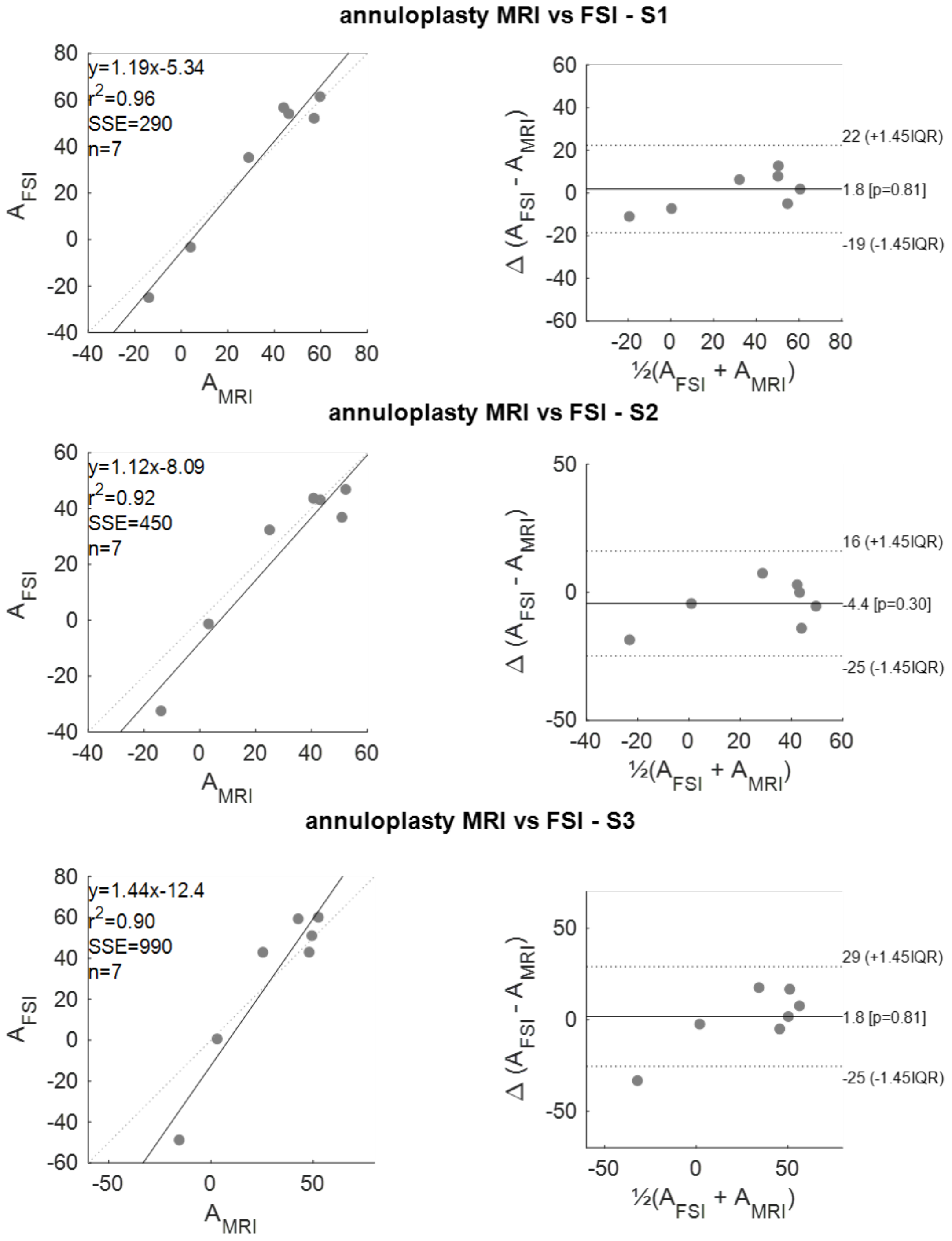


**Fig. S7.1** – Correlation plots and non-parametric Bland-Altman analyses between the MRI and FSI annuloplasty models.

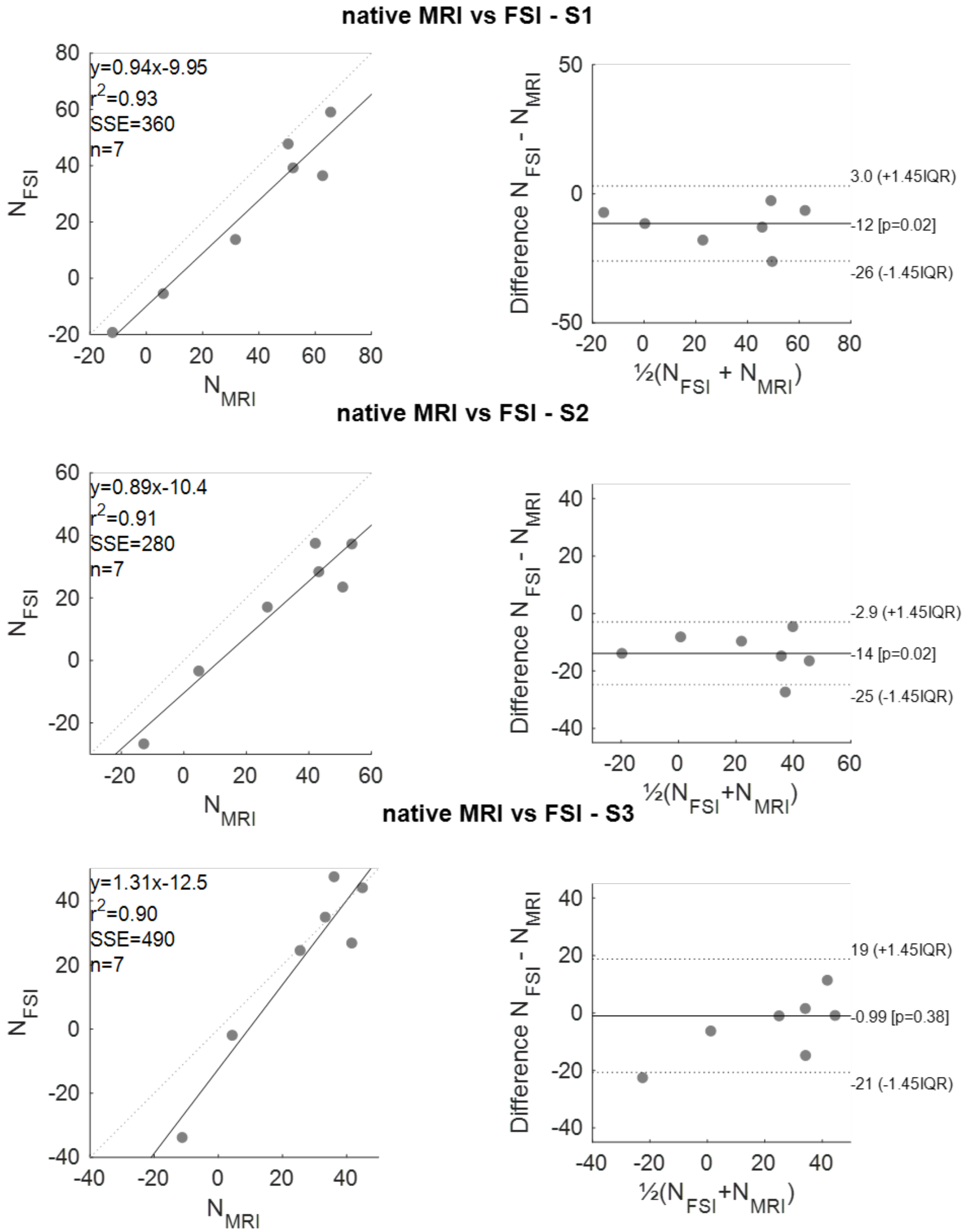


**Fig. S7.2 –** Correlation plot and Bland-Altman analysis between the MRI and FSI native models.

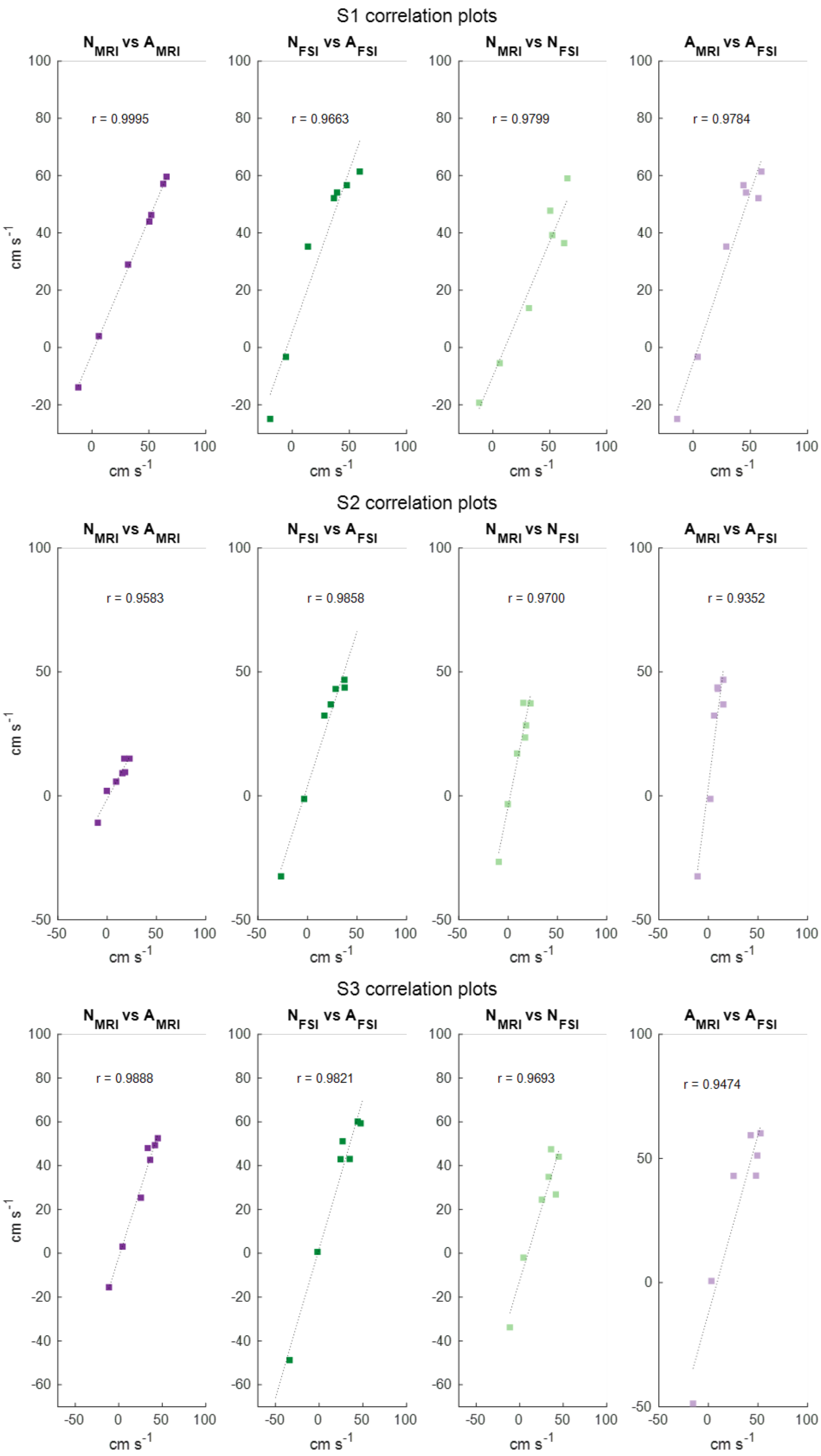


**Fig. S7.3** –Linear correlations plots describing the agreement between MRI and FSI methods, native (N) and annuloplasty (A) models.

[3] Formlabs. Formlabs Form 3B+, https://formlabs.com/it/3d-printers/form-3b/ n.d.
